# A signalling axis involving CNOT3, Aurora B and ERK promotes differentiation and survival of mesendodermal progenitor cells

**DOI:** 10.1101/756932

**Authors:** Moumita Sarkar, Matteo Martufi, Monica Roman-Trufero, Yi-Fang Wang, Chad Whilding, Dirk Dormann, Pierangela Sabbattini, Niall Dillon

**Affiliations:** Gene Regulation and Chromatin Group, Imperial College London, Hammersmith Hospital Campus, Du Cane Road, London W12 0NN, UK; Bioinformatics and Computing, Imperial College London, Hammersmith Hospital Campus, Du Cane Road, London W12 0NN, UK; Microscopy Facility, MRC London Institute of Medical Sciences, Imperial College London, Hammersmith Hospital Campus, Du Cane Road, London W12 0NN, UK

## Abstract

Mesendoderm cells are key intermediate progenitors that form at the early primitive streak (PrS) and give rise to mesoderm and endoderm in the gastrulating embryo. We have identified an interaction between CNOT3 and the cell cycle kinase Aurora B, which requires sequences in the NOT box domain of CNOT3, and regulates MAPK/ERK signalling during mesendoderm differentiation. Aurora B phosphorylates CNOT3 at two sites located close to a nuclear localization signal and promotes localization of CNOT3 to the nuclei of mouse ES cells (ESCs) and metastatic lung cancer cells. ESCs that have both sites mutated give rise to embryoid bodies that are largely devoid of mesoderm and endoderm and are composed mainly of ectoderm. The mutant ESCs are also compromised in their ability to differentiate into mesendoderm in response to FGF2, BMP4 and Wnt3. The double mutation affects interaction of CNOT3 with Aurora B and with ERK and reduces phosphorylation of ERK in response to FGF2, impacting on survival of the differentiated ME cells. Our results identify an adaptor function for CNOT3 that regulates a key pathway in embryogenesis and cancer.

## INTRODUCTION

Gastrulation in mammals and birds is initiated by differentiation of mesendoderm (ME) cells, which form at the primitive streak (PrS) as transient precursors expressing both mesodermal and endodermal markers in response to signalling from the visceral endoderm (VE)^1, 2^. Ingression of ME cells along the midline of the PrS during gastrulation is accompanied by differentiation into mesoderm and definitive endoderm and formation of the embryonic germ layers. Signalling ligands that have been shown to be involved in specifying ME differentiation and PrS formation in the early embryo include Wnt3, Activin, BMP4 and FGF2.^3–7^ In combination, these ligands can be used to induce ESC differentiation into ME, which is characterised by expression of the mesodermal marker Brachury and endodermal markers GATA4, GATA6 and FOXA2 (reviewed in ^8^).

The RAS/MEK/ERK signaling pathway regulates an extremely diverse range of cellular processes at all stages of development^9^ and has been shown to play a vital role in the formation of ME in mouse and human ESCs.^2, 10^ The diversity of functional roles for MAPK/ERK signalling implies the existence of multiple secondary levels of regulation that direct the signals towards their different targets. Adaptor proteins that bind more than one signalling component are one type of mechanism that can be used to regulate MAPK/ERK activity and channel it towards specific targets in a given cell type.^11, 12^

CNOT3 was first identified as a component of the CCR4-Not complex, which is involved in regulating transcription and RNA processing in the nucleus and mRNA turnover in the cytoplasm.^13^ Studies in ES cells have shown that CNOT3 acts in conjunction with the transcription factors cMYC and ZFX to form part of a transcriptional regulatory module, that binds to a number of gene promoters in ESCs^14^ and has been reported to inhibit differentiation of ESCs into extraembryonic lineages.^14, 15^ Knockout of *Cnot3* in the mouse leads to embryonic lethality at the blastocyst stage caused by loss of inner cell mass cells.^16^

In addition to its early embryonic functions, CNOT3 has been shown to have important roles in B lymphopoesis^17^, and maintenance of adipose tissues^18^ in adult mice. *Cnot3* has also been shown to have tumour suppressor and tumour promoting properties.^19–21^ Mutations that alter the protein sequence have been identified in a range of cancers, with the largest number observed in prostate and colon cancers.^22, 23^ The diverse biological and cellular roles of CNOT3 and the presence of the C-terminal specialist protein-protein interaction domain, the NOT box, and a C-terminal coiled-coil domain, linked by a long intrinsically disordered central region^24^ suggest that it could function as an adaptor that mediates cross-talk between different cellular pathways. Here we show that CNOT3 has the properties of an adaptor protein that links the cell cycle kinase Aurora B with the MAPK/ERK signalling pathway.

Aurora B and ERK interact with the NOT box domain of CNOT3 and phosphorylation of the protein by Aurora B at sites adjacent to a nuclear localisation signal (NLS) promotes localisation of CNOT3 to the nucleus and increases interaction between ERK and CNOT3. Mutation of the Aurora B target sites reduces the level of active phosphorylated ERK in ESCs, preventing efficient differentiation of ME in response to FGF. We also present evidence supporting involvement of Aurora B-mediated phosphorylation and nuclear localisation of CNOT3 in the tumorigenic effects of CNOT3.

## RESULTS

### Phosphorylation by Aurora B localises CNOT3 to ESC nuclei

The Aurora B cell cycle kinase phosphorylates multiple protein targets during mitosis and cytokinesis as part of the chromosomal passenger complex (reviewed in ^25^) and is also involved in regulating the G1 to S-phase transition.^26, 27^ Upregulation of Aurora B protein levels has been shown to be a marker for lymph node metastasis in lung, colon and breast cancers. ^28–30^ We initially identified a strong interaction between Aurora B and CNOT3 in a co-immunoprecipitation screen and analysis of primary resting and activated mouse B lymphocytes ^31^ (M. Martufi and N. Dillon, unpublished data). The known involvement of CNOT3 in ESC pluripotency^14^ led us to investigate whether Aurora B interacts with CNOT3 in ESCs and co-immunoprecipitation of ESC extracts with anti-Aurora B antibody confirmed that the two proteins interact in ESCs (Figure 1A).

**Figure 1:**
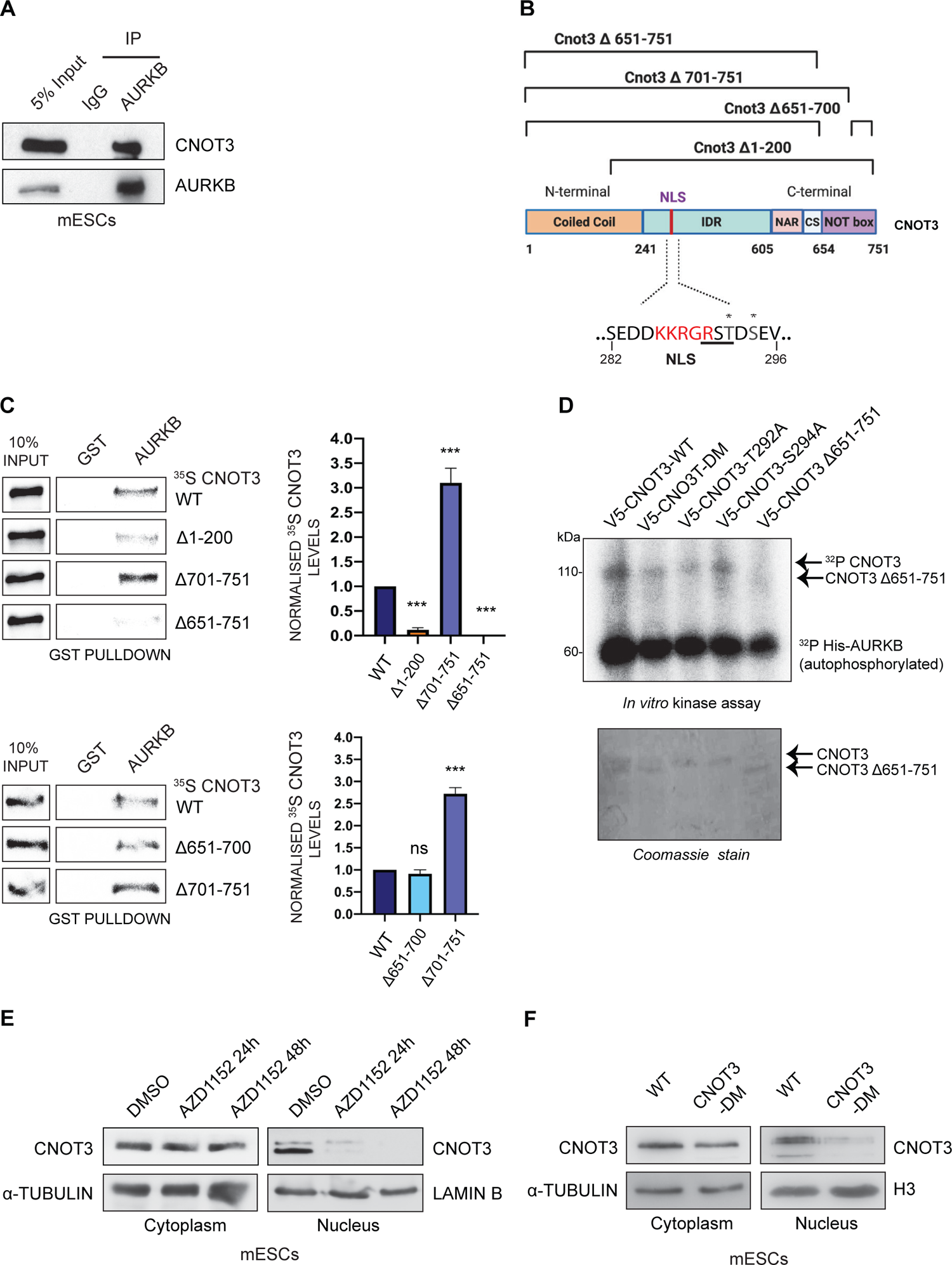
Phosphorylation by Aurora B is increases nuclear localisation of CNOT3. **A.** Representative immunoprecipitation (IP) carried out on cell extracts prepared from wild-type ESCs using anti-Aurora B (AURKB) antibody. Blots were probed with anti-CNOT3 antibody and anti-Aurora B antibody. IP with normal mouse IgG was used as a negative control. Input = 5% of the extract. **B.** Schematic representation of the organisation of the domains of CNOT3 ^24^ and the deletion mutants of *Cnot3* used in the GST pulldown assay. IDR = intrinsically disordered region; NAR = NOT1 anchor region; CS = connector sequence; NLS (red type) = nuclear localisation sequence. The Aurora B consensus phosphorylation site is underlined. **C.** *In vitro* GST pulldown assay using GST-Aurora B and ^35^S-labelled *in vitro* transcribed/translated wild-type (WT) CNOT3 and CNOT3 deletion mutants (as shown in B). Top and bottom left panels: Representative pulldown assays of GST-Aurora B and ^35^S labelled WT CNOT3 and CNOT3 deletion mutants. GST-only was used as a negative control for each pulldown. Input =10% of the respective labelled proteins. Right panels: Histograms show quantification of the band intensities normalized to input and relative to WT-CNOT3. Mean ± SEM; *n*=3 (unpaired *t*-test ***P<0.001). **D.** Top panel: *In vitro* kinase assay (top) using *γ*-^32^P-ATP carried out on purified V5-tagged *in* vitro transcribed/translated WT CNOT3, and single mutant constructs (CNOT3-T292A or CNOT3-S294A), double mutant (CNOT3-DM), and CNOT3 *Δ*651-751. The lower band is autophosphorylated Aurora B. Bottom panel: Coomassie stain showing the levels of the respective proteins. **E.** Representative immunoblot analysis of CNOT3 from cytoplasmic and nuclear extracts of WT ESCs, treated with Aurora B inhibitor AZD1152 for 24 hours and 48 hours respectively before harvesting. DMSO: vehicle control. Cytoplasmic loading control: *α*-Tubulin. Nuclear loading control: Lamin B. **F.** Representative immunoblot analysis of CNOT3 levels in cytoplasmic and nuclear extracts of wild-type (WT) and *Cnot3*-DM ESCs. Cytoplasmic loading control: *α*-Tubulin. Nuclear loading control: Histone H3.

Direct interaction of specific regions of CNOT3 with Aurora B was demonstrated by using GST-tagged Aurora B for *in vitro* pulldown of ^35^S-labelled wild-type and CNOT3 deletion mutant proteins (Figure 1B, 1C, Figure S1A). The results of the deletion analysis showed that deletion of the entire C-terminal NOT box domain (*Δ*651-751) results in near-complete loss of Aurora B binding.

Deletion of the N-terminal segment of the NOT box (*Δ*651-700) had no effect on binding of Aurora B, whereas deletion of the C-terminal region of the NOT box (*Δ*701-751) strongly increased the amount of interaction observed. (Figure 1C, bottom panels). The N-terminal region of CNOT3 (*Δ*1-200) was also required for full binding, indicating a complex interaction between Aurora B and CNOT3, with binding dependent on the NOT box domain, but with other regions involved in determining the level of interaction.

*In vitro* phosphorylation of CNOT3 by Aurora B, followed by mass spectrometry analysis, revealed phosphorylation at residues T292 and S294 in the CNOT3 protein (Figure S2A). T292 is located in a consensus Aurora B phosphorylation sequence (^290^R-S-T^292^) (Figure 1B). Phosphorylation of purified wild-type and mutant CNOT3 proteins by Aurora B in the presence of [*γ*-^32^P]-ATP showed reduced labelling of a double CNOT3-T292A/S294A mutant protein (Figure 1D). A reduction in labelling was also observed when T292 alone was mutated to A, but the effect was less pronounced when S294 was mutated on its own. Deletion of the NOT box region abolished labelling, indicating that docking of Aurora B on the NOT box is required for the kinase to phosphorylate CNOT3 (Figure 1D).

Inspection of publicly available high-throughput mass spectrometry data from human tissues^32^ revealed that the T292/S294 residues are part of a phosphorylation hotspot extending from residues 291-299 of CNOT3 (Figure S1B). The phosphorylation hotspot, which includes T292 and S294, is located adjacent to a sequence motif (^286^K-K-R-G-R^290^) (Figure 1B, Figure S1A), which has been shown to form part of a functional nuclear localisation sequence (NLS) in the *Toxoplasma gondii* GCN5-B histone acetyl transferase and the Influenza D virus nucleoprotein^33–35^, suggesting that phosphorylation of this region could be involved in nuclear localisation. To directly test this idea, ESCs were incubated for 24 and 48 hours with the specific Aurora B inhibitor AZD1152.

Comparison of the levels of CNOT3 in the nuclear and cytoplasmic fractions after 24 hours incubation with AZD1152 showed a substantial reduction in the amount of CNOT3 in the nucleus (Figure 1E) and the level was further reduced after 48 hours of treatment. The level of CNOT3 in the cytoplasm was largely unaffected (Figure 1E) and inhibition of Aurora B did not significantly affect the cell cycle of the ESCs (Figure S2C and S2E). These results provide evidence that phosphorylation of CNOT3 by Aurora B is involved in specifying localisation of CNOT3 to the ESC nucleus. We confirmed this by transfecting V5-tagged CNOT3 expression constructs carrying mutations of either T292 or S294, or of both residues, to alanine, into HEK293 cells. The results showed reduction of nuclear CNOT3 for each of the single mutants and the double mutation, indicating that both Aurora B target sites contribute to nuclear localisation in this assay (Figure S2F).

CRISPR/Cas9 mutagenesis was used to simultaneously mutate the T292 and S294 residues to alanine in mouse ESCs (Figure S2B). Generation of three ESC clones that were homozygous for the T292A/S294A double mutation (*Cnot3*-DM) was confirmed by sequencing (Figure S2B). The mutant ESCs grew normally and had the characteristic cell cycle profile of ESCs (Figure S2D and S2E). Analysis of nuclear CNOT3 levels showed a substantial reduction in the amount of CNOT3 in the nuclei of *Cnot3*-DM ESCs compared with WT cells (Figure 1F). Cytoplasmic levels of CNOT3 were largely unchanged in the mutant cells.

### Effect of mutating CNOT3-T292/S294 on embryoid body formation

To test whether the absence of phosphorylation of CNOT3-T292/S294 affects ESC pluripotency and differentiation potential, wild-type and *Cnot3*-DM ESCs were cultured under conditions that promoted the formation of embryoid bodies (EBs) (see Methods for details). The results show that the double *Cnot3* mutation (*Cnot3*-DM) resulted in a 40% reduction in EB numbers after 10 days in culture and the average size of the EBs was reduced by 40-50% (Figure 2A and 2B). Staining of the EBs for the lineage markers Brachyury (mesoderm), FOXA2 (Endoderm) and Nestin (Ectoderm) showed that all three markers were strongly expressed in EBs formed from wild-type ESCs, with Brachyury giving broad staining of the central and peripheral regions of the EBs, whereas FOXA2 staining was more restricted to the central regions (Figure 2C). The ectodermal Nestin staining was broadly distributed in the EBs but was strongest at the periphery. In contrast, the *Cnot3*-DM EBs showed very low staining for Brachyury and FOXA2 and strong central and peripheral staining for Nestin (Figure 2C). This was confirmed by staining for the ectodermal marker OTX2 (Figure S3A). The observation that the mutant EBs were predominantly composed of ectoderm suggests that blocking phosphorylation of CNOT3-T292/S294 interferes with germ layer formation.

**Figure 2:**
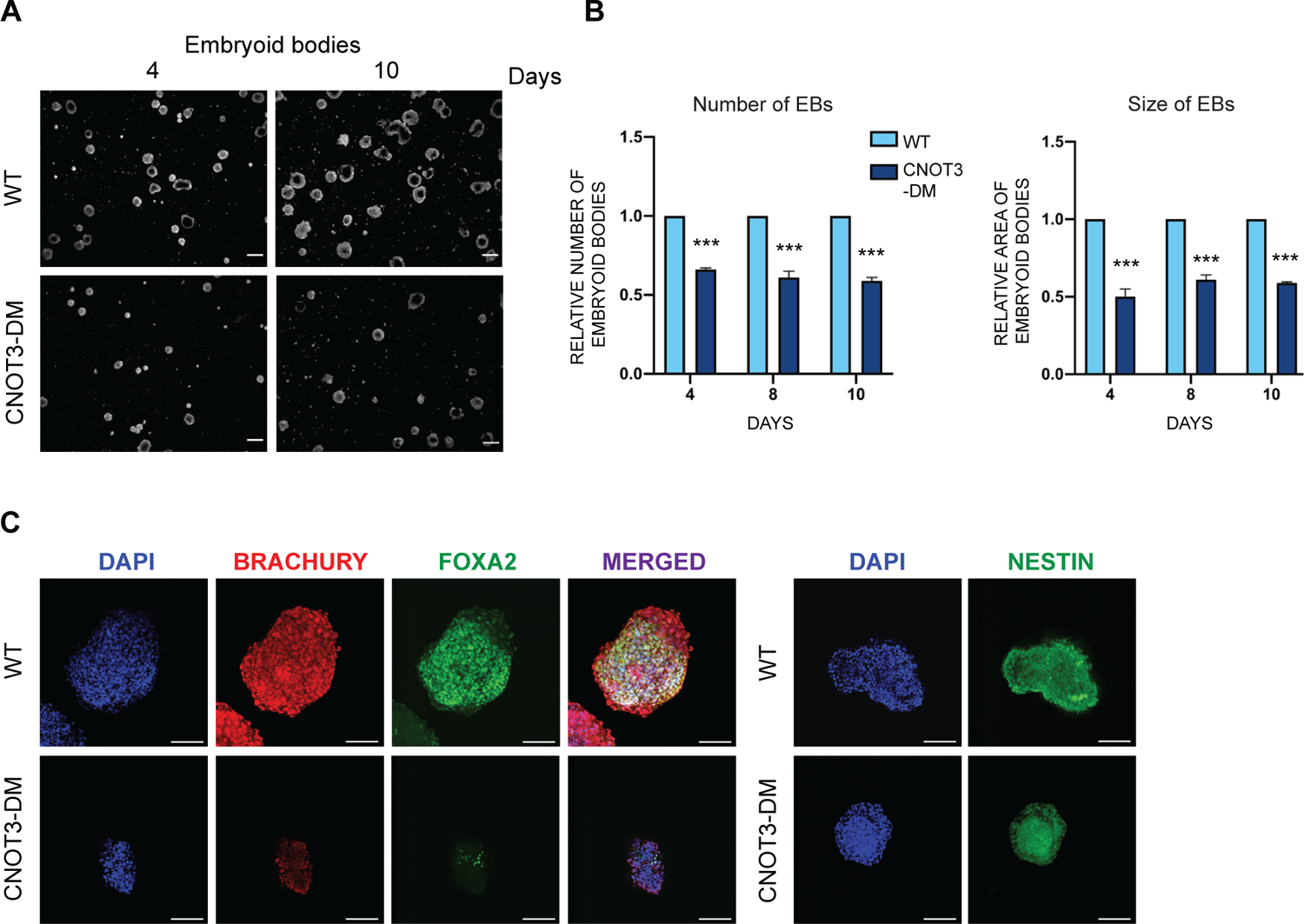
Mutation of CNOT3-T292A/S294 affects EB germ layer formation. **A.** Representative phase-contrast images of embryoid bodies (EBs) formed from wild-type (WT) and *Cnot3*-DM ESCs after 4 and 10 days. Scale bar = 100 μm. **B.** Histograms show the number (left panel) and size (right panel) of EBs formed from *Cnot3*-DM ESCs relative to WT cells at 4, 8 and 10 days respectively. Mean ± SEM; *t*-test ***P<0.001 *n*=3. **C.** Left panel: Representative confocal images of EBs derived from WT and *Cnot3*-DM ESCs stained with the mesodermal marker Brachury (*red*) and endodermal marker FOXA2 (green) at 8 days (scale bar = 100 μm). Right panel: staining of WT and *Cnot3*-DM EBs for ectodermal marker nestin (green) (scale bar = 100 μm). Nuclei were stained with DAPI. Staining for an additional ectodermal marker OTX2 is shown in Supplementary Figure 3A.

### CNOT3-T292/S294 phosphorylation promotes ME differentiation and survival

Transitioning of cells through an intermediate ME progenitor cell stage at the PrS ^36–38^, has been shown to depend on Nodal/Activin, Bmp, Wnt and FGF signaling pathways. ^2, 3, 10, 39–41^ We carried out differentiations of wild-type and *Cnot3*-DM ESCs in a defined medium containing BMP4 and FGF2 and different combinations of Activin, Wnt and the GSK3*β* inhibitor CHIR99021, which stabilises *β*-catenin, bypassing Wnt activation ^42^ (Figure 3A-3C) (see Methods). When applied to wild-type cells, all of the combinations that included BMP4 and FGF2, and either Wnt, Activin or CHIR99021 gave similar numbers of differentiated cells (Figure 3A, 3B) with a high proportion staining for Brachyury and FOXA2 (Figure 3C). Incubation with FGF2 and BMP4 alone also promoted ME differentiation of wild-type ESCs, but the number of differentiated cells was lower. The mesendodermal identity of the cells was confirmed by analysis of mRNA levels for mesodermal and endodermal lineage markers by qRT-PCR (Figure S3C). Analysis of the *Cnot3*-DM cells showed that they gave significantly reduced numbers of differentiated cells for all ligand combinations tested, (Figure 3A and 3B), with the largest reductions relative to wild-type observed with Wnt + FGF2 + BMP4 (80%) and FGF2 + BMP4 (65%). Incubation of wild-type cells with activin on its own gave reduced survival compared with incubation with FGF2 + BMP4, whereas survival of the mutant cells was similar under both conditions (Figure S3B). In contrast, cells that were incubated with Wnt in the absence of the other signalling ligands showed a reduction of around 10-fold in survival of wild-type cells and an even greater reduction for the *Cnot3*-DM cells (Figure S3B). This result is consistent with FGF2 + BMP4 have a major role in promoting survival of the differentiating mesendodermal cells.

**Figure 3:**
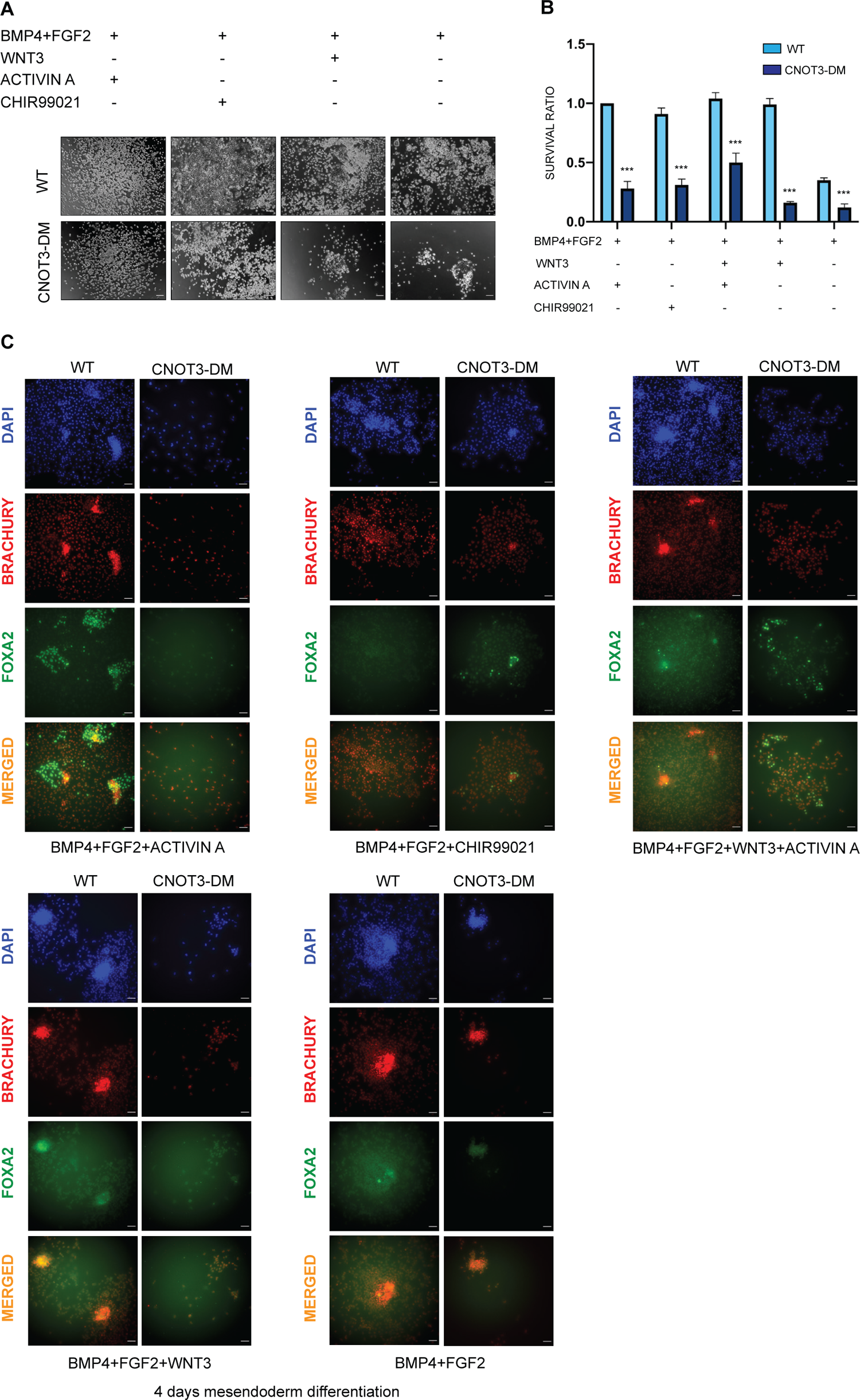
CNOT3 phosphorylation promotes efficient differentiation of ME. **A**. Representative phase-contrast images showing the efficiency of mesendoderm (ME) differentiation of wild-type (WT) and *Cnot3*-DM ESCs over 4 days in the presence of various combinations (as indicated above the images) of Activin A, CHIR99021, Wnt3 and BMP4 + FGF2. Scale bar = 100 μm. **B**. Histogram shows survival of WT and *Cnot3*-DM ESCs treated with different combinations of the ligands used in (A) over 4 days of ME differentiation. Activin A, CHIR99021 or Wnt3 was added after two days of BMP4+FGF2-induced differentiation and the cell survival was determined using the WST-1 reagent on day 4 of the differentiation. Survival ratios (mean ± SEM) were calculated relative to the values obtained for WT cells treated with Activin A+ BMP4+FGF2, which was assigned a value of 1. Significance was calculated between WT and CNOT3-DM for each combination using unpaired *t*-test ***P<0.001, *n*=3. Analysis of cell survival in the presence of Activin only or Wnt only is shown Figure S4D. **C.** Representative IF images of mesodermal marker Brachury (*red*) and endodermal marker FOXA2 (green) following ME differentiation of WT and *Cnot3*-DM ESCs for 4 days induced by combinations of BMP4 + FGF2, Activin A, CHIR99021 and Wnt3 (as indicated). Merged red and green images show the ME cells. Nuclei were stained with DAPI; scale bar = 100 μm.

A time-course from day 2 to day 8 of ME differentiation in the presence of BMP4 + FGF2 only, showed reduced numbers of *Cnot3*-DM cells from day 2 onwards (Figure 4A, 4B). Staining for additional mesodermal and endodermal lineage markers (SMA: mesoderm; GATA4: endoderm) after 4 and 8 days of incubation with confirmed ME differentiation (Figure S4A and S4B). Analysis of the differentiation capacity of the other two *Cnot3*-T292A/S294A mutant clones that were generated by the CRISPR/CAS9 targeting showed a similar failure to expand and proliferate in the mesendoderm differentiation medium (Figure S5B), confirming that the effect was caused by the double mutation. In contrast, differentiation of the *Cnot*3-DM cells directly into mesoderm, endoderm and ectoderm was largely unaffected with only endoderm differentiation induced by FGF2 and retinoic acid showing any reduction in the mutant cells (Figure S5A). These findings support the idea that the regulatory role of CNOT3-T292/S294 phosphorylation in ME differentiation and survival is linked to specific signalling pathways and suggest that FGF signalling is one of these pathways.

**Figure 4:**
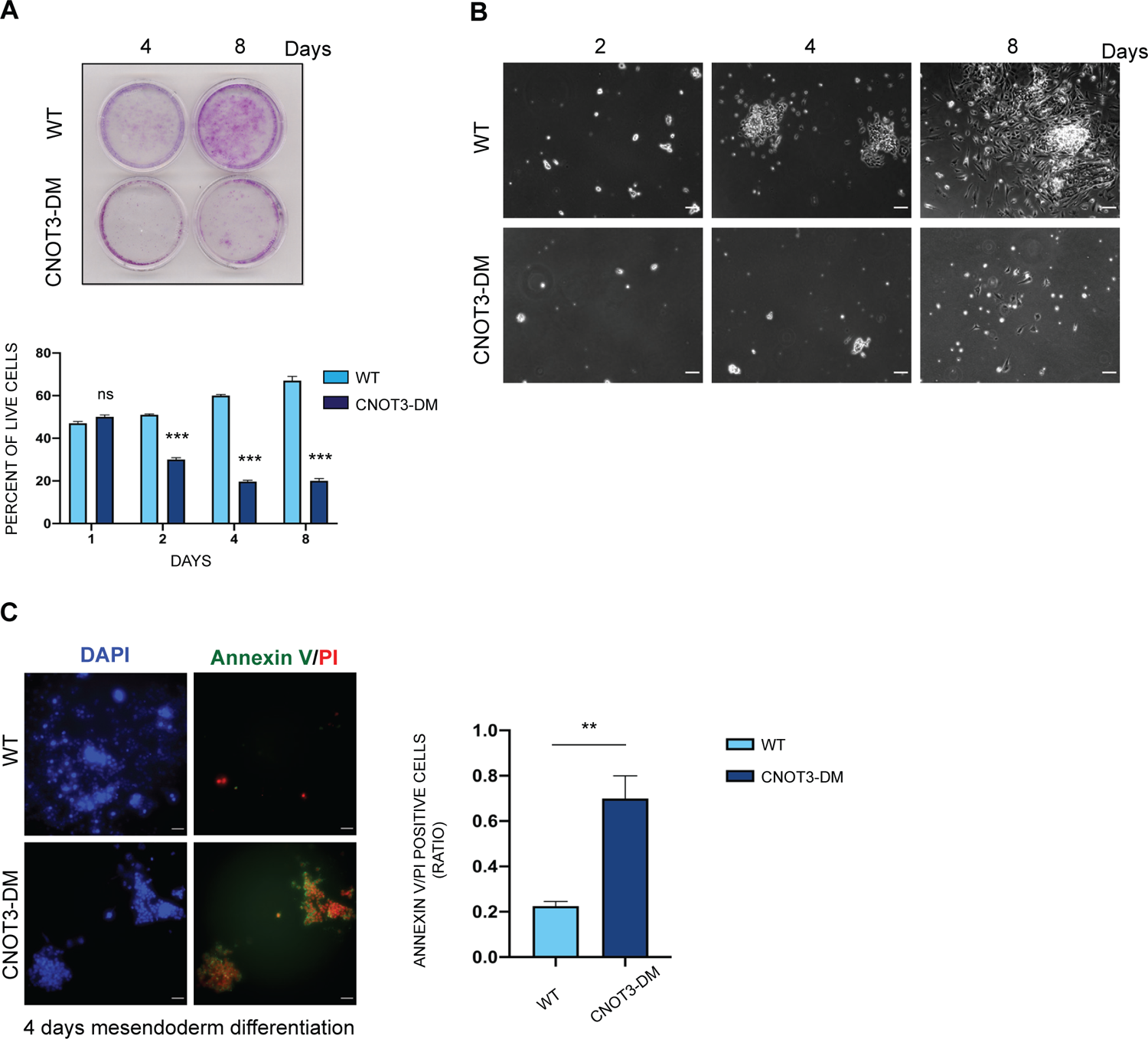
Phosphorylation of CNOT3 prevents apoptosis of differentiating ME cells. **A**. Top panel: Crystal violet staining of cells after 4 days BMP4 + FGF2-induced mesendoderm differentiation of wild-type (WT) and *Cnot3*-DM ESCs after 4 and 8 days. Bottom panel: At the indicated time-points, live cells were counted based on trypan blue exclusion. Histogram shows the percentage of live cells at different time points of differentiation. Mean ± SEM; unpaired *t*-test between WT and *Cnot3*-DM for each time-point, ***P<0.001 *n*=3. Differentions of two other independently generated clones of *Cnot3*-T292A/S294A mutant ESCs are shown in Figure S3A. **B**. Representative phase-contrast images showing the efficiency of BMP4 + FGF2 induced mesendoderm differentiation of WT and *Cnot3*-DM ESCs for 2, 4, and 8 days. Scale bar: 100 μm. See also Supplementary Video 1 for a time-lapse analysis of the differentiations. **C**. IF analysis of apoptotic markers Annexin V (*green*) and PI (red) in cells differentiated for 4 days with BMP4 + FGF2. Merged images show the Annexin V and PI stained apoptotic cells. Nuclei were stained with DAPI; scale bar = 100 μm. Histogram represents ratio of Annexin V/PI positive cells relative to total number of cells. Apoptotic cells were counted from ten randomly chosen fields for each biological replicate. Mean ± SEM; *t*-test **P<0.01 *n*=3.

The conclusion that proliferation and survival of ME cells is strongly affected by the double mutation was further reinforced by the results of a time-lapse analysis of the differentiating wild-type and mutant cells between 4 and 8 days of ME differentiation (Supplementary Video 1). The time-lapse analysis showed an explosive proliferation of the wild-type cells, whereas the *Cnot3*-DM cells failed to expand and appeared to undergo high rates of cell death following cell division. The cell cycle profiles of the wild-type and mutant cells after 4 days of differentiation were similar, implying that the major effect of the mutation was on survival of differentiating ME cells (Figure S5C).

Susceptibility of the *Cnot3*-DM cells to apoptosis during ME differentiation in response to FGF2 and BMP4 was directly assessed by staining wild-type and mutant cells with the apoptotic marker Annexin V and also by measuring propidium iodide (PI) uptake after 4 days of differentiation. The wild-type cells showed almost no staining for either cell death indicator whereas the 4-day differentiated mutant cells were strongly stained for PI and Annexin V, indicating that the failure of the mutant cells to differentiate was associated with high rates of apoptosis (Figure 4C). This result suggests a functional relationship between CNOT3 phosphorylation and survival signals mediated by incubation with FGF2 and BMP4.

Further support for the conclusion that BMP4 + FGF2 promote survival of the differentiating mesendoderm cells was provided by a transcriptomic comparison of the wild-type and *Cnot3*-DM cells using RNA-seq (Figure S6A and S6B). The analysis showed that 153 genes were significantly upregulated and 155 were downregulated in the mutant cells (Supplementary Tables 3A and 3B). Gene Ontology and GSEA analysis revealed that the mutation resulted in increased expression of genes that are associated with cell death and apoptosis (Figure S6C and S6E). In addition, genes involved in cell proliferation and the MAPK cascade were downregulated in the *Cnot3*-DM cells (Figure S6D). The RNA-seq analysis also showed that the level of mRNA for FGFR1 which is the major FGFR that promotes ME differentiation ^43–45^ was unchanged in the *Cnot3*-DM cells.

### Mutation of CNOT3-T292/S294 reduces ERK phosphorylation

ERK1/2 phosphorylation by MEK is one of the key events in the response to stimulation by FGF. ^9^ ERK has been reported to upregulate Aurora B expression in melanoma cells ^46^ and phosphorylated ERK plays important roles in cell survival. ^47^ Involvement of the FGF/MEK/ERK pathway in signalling cell survival during ME differentiation was supported by the observation that treatment of the cells with FGFR and MEK inhibitors caused a dramatic reduction in cell numbers (Figure 5A).

**Figure 5:**
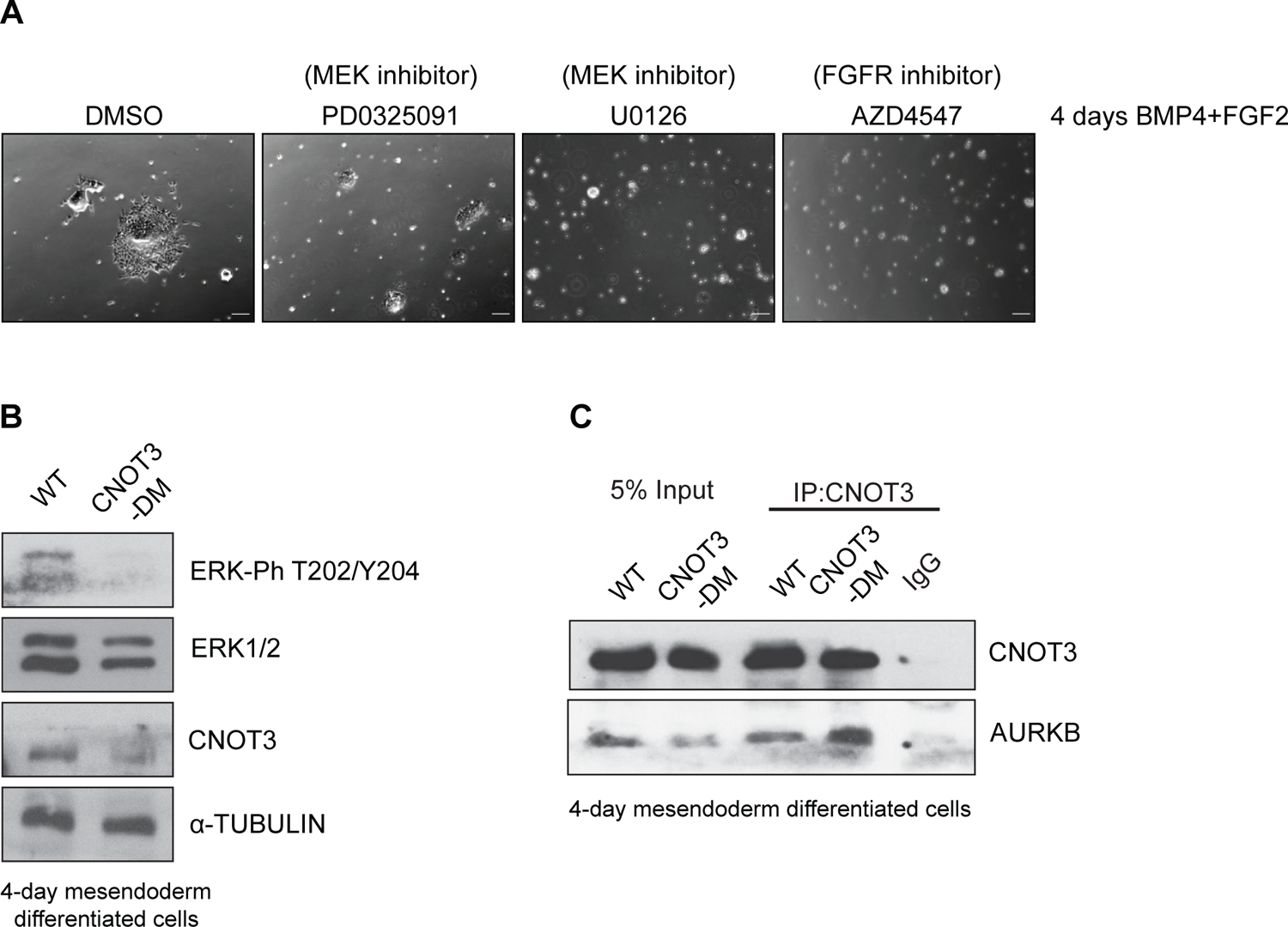
The CNOT3 double mutation reduces ERK phosphorylation during ME differentiation. **A**. Phase-contrast images showing the effects of MEK inhibitors (PD0325091 and U0126), which abolish ERK activity, and the FGFR inhibitor AZD4547, on BMP4 + FGF2 induced ME differentiation of wild-type (WT) ESCs. Differentiations were carried out for 4 days. Each inhibitor was added after two days of differentiation. Vehicle = DMSO. Scale bar = 100 μm. **B**. Representative immunoblot analysis of the indicated proteins carried out on cell extracts from WT and *Cnot3*-DM ESCs subjected to 4 days of BMP4 + FGF2 induced differentiation into ME. Loading control: *α*-Tubulin. **C**. Co-IP was carried out with anti-CNOT3 antibody on cell extracts prepared from WT and *Cnot3*-DM ESCs differentiated as in (B). Co-immunoprecipitated proteins were immunoblotted and probed with anti-CNOT3 and anti-Auorora B. Negative control: rabbit IgG. Input = 5% of extracts.

To assess the effect of the *Cnot3* T292A/S294A double mutation on ERK phosphorylation, extracts from wild-type and mutant ESCs that had been differentiated with FGF2 and BMP4 for 4 days were analysed by western blotting using an antibody that recognised the phosphorylated ERK TEY motif ^9^ and an antibody against total ERK. The results showed a strong reduction in the level of phosphorylated ERK in the *Cnot3*-DM cells compared to wild-type (Figure 5B).

Levels of phosphorylated ERK and of total ERK1/2 were unaffected in undifferentiated *Cnot3*-DM ESCs (Figure S7A). Aurora B and CNOT3 continue to interact in differentiating ME cells, as shown by immunoprecipitations carried out with anti-CNOT3 antibody on extracts from the 4-day differentiated wild-type and Cnot3-DM cells (Figure 5C). Levels of total CNOT3 were reduced in differentiated *Cnot3*-DM cells (Figure 5B), suggesting that phosphorylation by Aurora B stabilises wild-type CNOT3.

### Altered interaction of mutant CNOT3 with Aurora B and ERK

The proximity ligation assay (PLA) was used to analyse interactions between CNOT3, Aurora B and ERK at the single-cell level. PLA analysis with mouse anti-Aurora B and rabbit anti-CNOT3 antibodies after 4 days of differentiation of wild-type ESCs with BMP4 + FGF2 showed a high level of association between CNOT3 and Aurora B (Figure 6A). The number of interaction foci in the nuclei of wild-type cells was around half the number observed in the cytoplasm. Analysis of the *Cnot3*-DM cells showed a striking and unexpected increase of approximately 3-fold in the interaction between CNOT3 and Aurora B in the mutant cells compared to the wild-type cells and an increase in the proportion of interaction foci observed in the nucleus relative to the cytoplasm relative to wild type cells (Figure 6A). Overall, these results confirm the strong interaction between CNOT3 and Aurora B that was observed by co-IP of extracts from undifferentiated ESCs and after 4 days of ME differentiation. The differences between the interaction patterns observed in the wild-type and Cnot3-T292A/S294A mutant cells suggest a dynamic cycle of phosphorylation of T292/S294 and subsequent release of Aurora B in differentiating wild-type ME cells.

**Figure 6:**
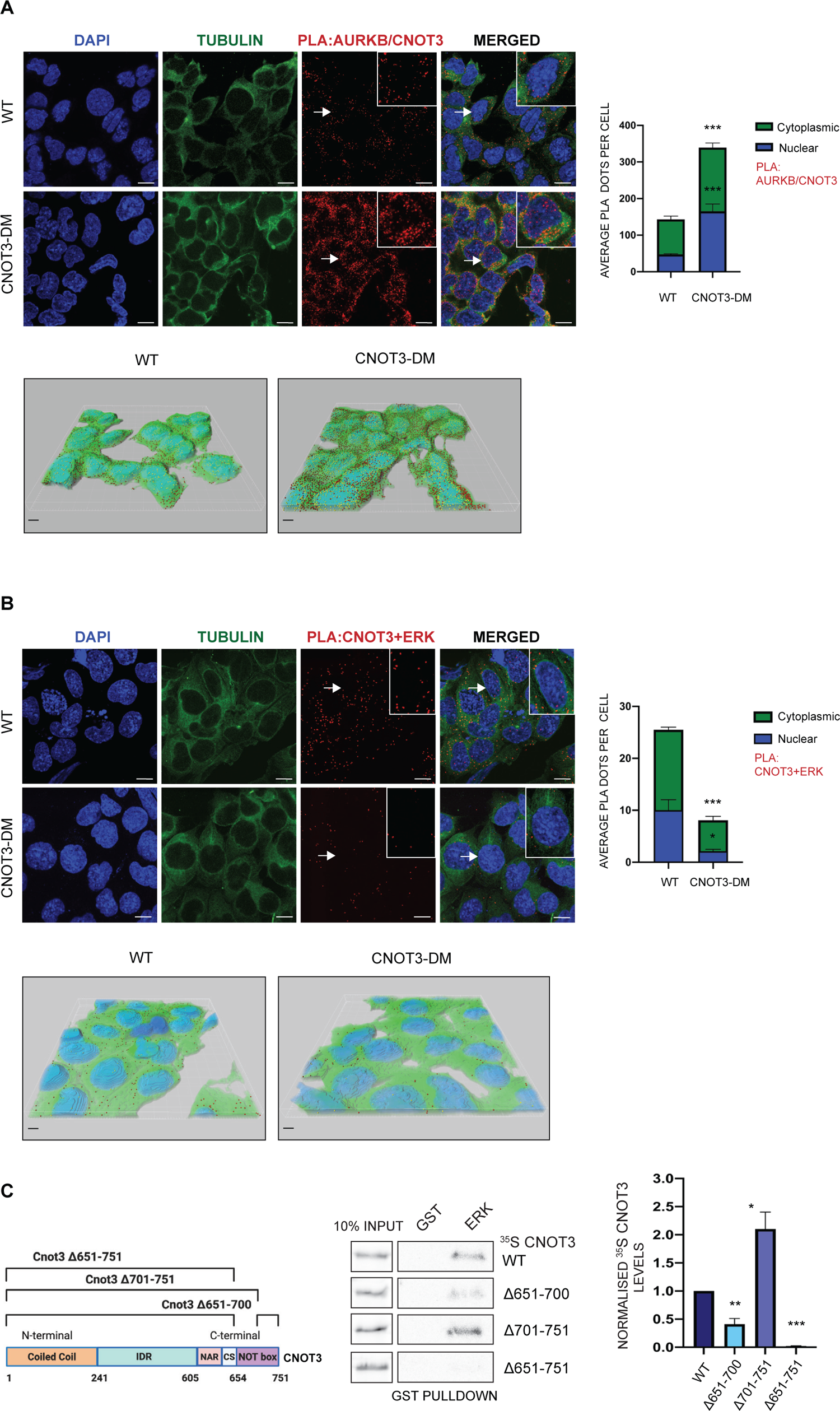
Phosphorylation of CNOT3 alters its interaction with Aurora B and ERK in ME cells. **A**. and **B.** PLA was used to detect interaction between endogenous Aurora B (AURKB) and CNOT3 (A) and CNOT3 and ERK (B) after 4 days of BMP4 + FGF2 induced differentiation of ME cells from wild-type (WT) and *Cnot3-*DM ESCs. Top panels: Red dots represent positive PLA signals. Nuclei were stained with DAPI and cytoplasm with anti-tubulin. Boxed areas show an enlarged image of a cell (indicated by the arrows) from each merged panel. scale bars =10 μm. Bottom panels: Three-dimensional images of the cells were digitally reconstructed, showing the distribution of PLA dots in the nucleus (yellow) and cytoplasm (red). scale bars = 5 μm.. PLA dots were quantified from randomly chosen fields from at least 50 cells for each biological replicate. Histograms represent average number of interactions per cell (dots/cell) in the nucleus (blue) and cytoplasm (green). Mean ± SEM; unpaired *t*-test ***P<0.001 (A) *n*=3 (B) *n*=2. Single antibody controls and PLA of CNOT3 and ERK using a second set of antibodies are shown in Figure S7C. **C.** *In vitro* GST pulldown assay using GST-ERK1 and *in vitro* transcribed/translated wild-type (WT) CNOT3 and CNOT3 deletion mutants labelled with ^35^S-methionine. Left panel: Schematic representation of the deletion mutants of *Cnot3*. Middle panel: Representative *in vitro* GST pulldown assay of GST-ERK1 and ^35^S-labelled WT CNOT3 and CNOT3 deletion mutants. GST-only was used as a negative control for each pulldown. Input =10% of the respective labelled proteins. Right panel: histogram shows quantification of the band intensities normalized to input and relative to WT CNOT3. Mean ± SEM; *n*=3 (unpaired *t*-test *P<0.05, **P<0.01, ***P<0.001).

PLA of the interaction between CNOT3 and ERK showed a substantial level of interaction in wild-type 4-day differentiated cells that was broadly distributed between cytoplasmic and nuclear compartments (Figure 6B). The *Cnot3*-DM cells showed a 3-fold reduction in the number of interaction foci in the cytoplasm and a 5-fold reduction in the nucleus. These results, which were confirmed with a second, separate anti-CNOT3/anti-ERK antibody pair (Figure S7B), provide strong evidence that ERK interacts with CNOT3 and that this interaction is promoted by Aurora B-mediated phosphorylation of CNOT3-T292/S294. The observation that mutation of the Aurora B phosphorylation sites reduced phosphorylation of ERK suggests that the interaction between ERK and phosphorylated CNOT3 promotes ERK phosphorylation or stabilises phosphorylated ERK, thereby enhancing Ras/MEK/ERK signalling.

*In* vitro pulldown of ^35^S-labelled CNOT3 with GST-ERK showed that ERK interacts directly with CNOT3 (Figure 6C). Analysis of CNOT3 deletion mutants showed some similarities to the binding of Aurora B, with deletion of the NOT box abolishing binding and deletion of the C-terminal region enhancing the interaction with GST-ERK. However, the effects of the CNOT3 deletions also showed some differences from the effects that we observed on Aurora B binding (Figure 1C), with the *Δ*651-700 deletion mutant giving a 50% reduction in ERK binding whereas the same mutation did not show a reduction in binding of Aurora B. This implies that there are differences in the contacts that Aurora B and ERK make with the CNOT3 NOT box region.

### Phosphorylation alters localisation of CNOT3 in cancer cells

Mutations in the coding region of human *Cnot3* have been observed in a number of cancers ^22^, with genetic analysis providing evidence that *Cnot3* mutations can have tumour suppressor ^19^ and tumour promoting ^48^ effects. High levels of nuclear CNOT3 have also been observed in an aggressive colorectal cancer cell line ^21^. To test whether Aurora B might be promoting nuclear localisation and EMT/MET in cancer cells, H1299 and A549 non-small cell lung cancer (NSCLC) cells were stained with anti-CNOT3 antibody. H1299 was originally derived from a lymph node metastasis and A549 from a lung adenocarcinoma. The results of the immunofluorescence analysis (Figure 7A) showed that nuclear CNOT3 levels were approximately 3-fold higher than the cytoplasmic levels in H1299 cells, whereas the difference was 1.6-fold in A549 cells.

**Figure 7:**
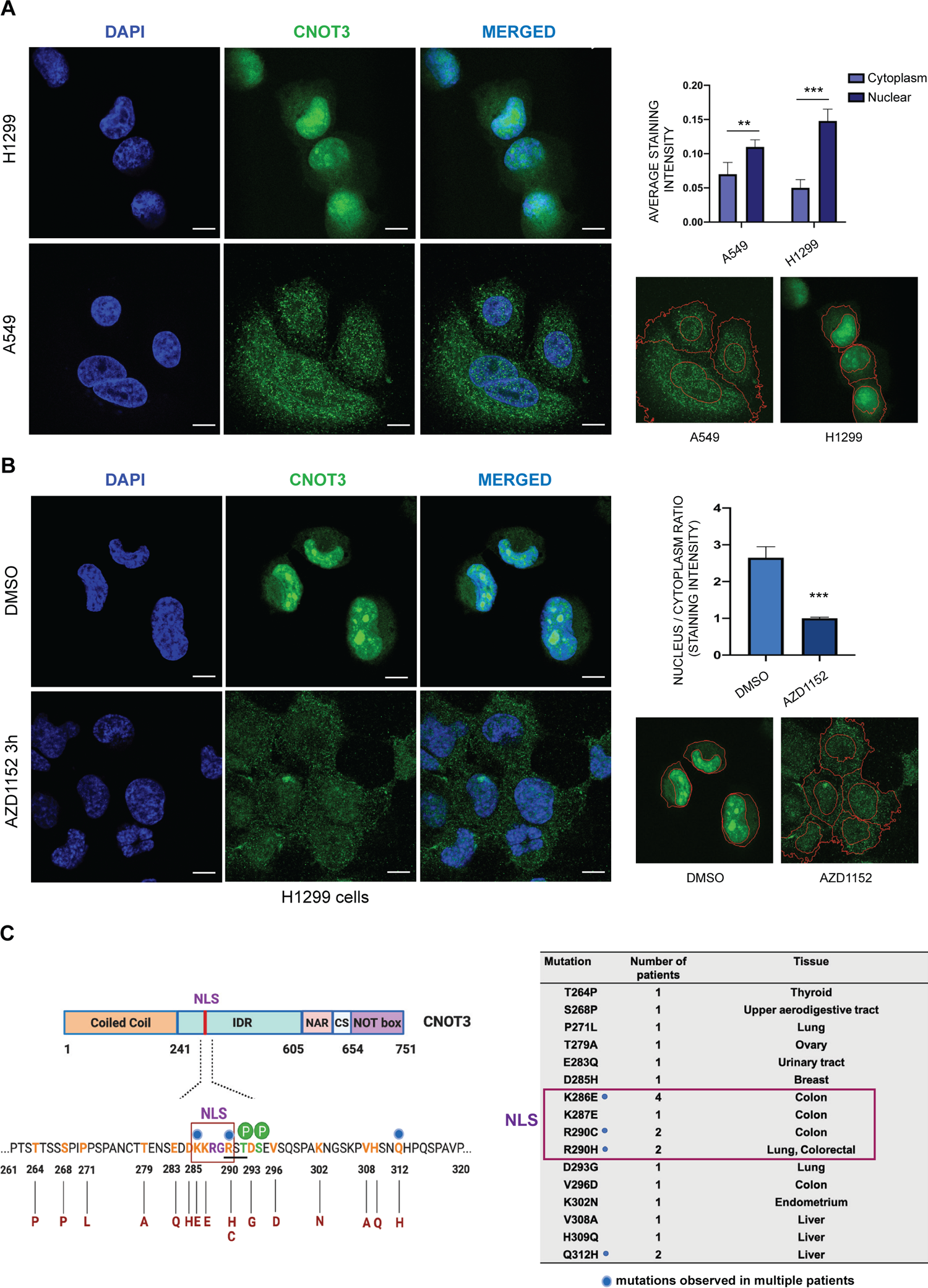
Phosphorylation by Aurora B promotes nuclear localisation of CNOT3 in a metastatic NSCLC cell line. **A.** Comparison of nuclear and cytoplasmic localisation of CNOT3 in two NSCLC cell lines. The H1299 line was derived from a lymph node metastasis and the A549 line from a lung adenocarcinoma. Left panel: H1299 cells (top) and A549 cells (bottom) immunostained with anti-CNOT3 and counterstained with DAPI (scale bar: 10 μm). Right panel: Histogram shows digital quantification of the average intensities of the nuclear and cytoplasmic staining for CNOT3 in the H1299 and A549 cells. Mean ± SEM; *n*=3 (unpaired *t*-test **P<0.01, ***P<0.001). Images (bottom) show representative fields depicting segmentation of the cells and of the nuclei used for the quantification. Quantification was performed on at least 30 cells from randomly chosen fields for each biological replicate. **B.** Effect of the Aurora B inhibitor AZD1152 on nuclear and cytoplasmic localization of CNOT3 in H1299 cells. Left panel panel: H1299 cells were incubated for 3 hours with AZD1152 (bottom) or with vehicle (DMSO, top) and were then immunostained as in (A) (scale bar = 10 μm). Right panel: Histogram shows quantification of the average ratio of nuclear to cytoplasmic staining for CNOT3 in the AZD1152 and vehicle-treated cells. Mean ± SEM; *n*=3 (unpaired *t*-test ***P<0.001). Images (bottom) show representative fields as in (A). Quantification was performed on at least 30 cells from randomly chosen fields for each biological replicate. **C.** Missense mutations described in human cancer patients in the region of CNOT3 spanning residues 261 to 320. Blue spheres indicate mutations that have been observed in more than one patient. The nuclear localisation signal (NLS) is indicated by red rectangles in the left and right panels and the Aurora B consensus is underlined in the left panel. P = phosphorylation sites.

Involvement of Aurora B phosphorylation in the H1299-specific localisation of CNOT3 was investigated by treating H1299 cells for 3 hours with the Aurora B inhibitor AZD1152 (Figure 7B). The incubation time was optimised to avoid disruption of the cell cycle. The results showed a dramatic effect of Aurora B inhibition on CNOT3 localisation in the H1299 cells with the nuclear levels of CNOT3 in the inhibitor-treated cells reduced by 3-fold relative to the cytoplasmic level (Figure 7B).

The involvement of the NLS and adjacent Aurora B target sites in cancer is further supported by the recent identification of a *Cnot3*-K286E mutation in 11% (4/37) of a cohort of pre-malignant adenomas from that were sequenced from Familial Adenomatous Polyposis (FAP) patients ^48^ (Figure 7C). *Cnot3*-K286 is the first residue of the KKRGR nuclear localisation sequence that is located adjacent to the sites of Aurora B phosphorylation in mouse ESCs. Mutations that affect the first residue of the Aurora B consensus (*Cnot3*-R290C and -R290H) have also been described in one lung and three colorectal cancers. ^22, 23, 49^ Overall, a total of nine patients with cancer or pre-cancerous lesions have been shown to have mutations in the NLS (Figure 7C). This clustering of mutations provides strong evidence of a role for this region in oncogenic progression, particularly in colorectal cancers.

## DISCUSSION

The epithelial to mesenchymal transition (EMT) that leads to mesendoderm differentiation and gastrulation is a critical stage in mammalian embryogenesis. Our results identify a role for CNOT3 in regulating the signalling pathways that promote mesendoderm differentiation in the early embryo. The effect occurs through a direct interaction between CNOT3 and the Aurora B kinase, that is completely dependent on the NOT box domain of CNOT3, and results in Aurora B-mediated phosphorylation of specific sites adjacent to an NLS within the CNOT3 protein and in increased localization of CNOT3 to the nucleus. The involvement of the NLS-adjacent Aurora B phosphorylation sites in the CNOT3 protein in germ layer formation was dramatically illustrated by the observation that mutation of the two Aurora B target residues blocked mesoderm and endoderm differentiation in embryoid bodies formed from the mutant cells. *In vitro* differentiation of the mutant ESCs in a defined medium highlighted the importance of CNOT3 phosphorylation in FGF-driven differentiation and survival of ME progenitor cells and suggested that CNOT3 could also be involved in the cross-talk that is known to occur between the RAS/MEK/ERK and Wnt signalling pathways.^50, 51^

Analysis of ERK phosphorylation showed that ERK activation is affected by the *Cnot3* double mutation with a strong reduction in the levels of phosphorylated ERK observed in the mutant cells. This finding provides a direct mechanistic explanation for the increased apoptosis observed in the *Cnot3*-DM cells when differentiation is induced using FGF2 and BMP4. Phosphorylation of ERK by MEK is followed by rapid import of phospho-ERK into the nucleus, which occurs through a variety of mechanisms. ^47^ The level of phospho-ERK in the nuclear compartment is known to be strongly dependent on binding of ERK to stabilising proteins in the cytoplasm and in the nucleus. ^47, 52, 53^ The results of the *in vitro* GST pulldown and *in vivo* PLA analyses indicate that ERK interacts with CNOT3 via the NOT box domain in differentiating mesendodermal cells and that this interaction is strongly reduced in both the cytoplasmic and nuclear compartments of the Cnot3-DM cells. The observation that the double mutation increases interaction with Aurora B and reduces the CNOT3-ERK interaction, suggests a model (illustrated schematically in Figure 8) where the two kinases interact sequentially with CNOT3, with Aurora B first interacting with the CNOT3 NOT-box, followed by phosphorylation of CNOT3 at the NLS and release of the bound Aurora B, allowing ERK to bind to the NOT box. Our finding that Aurora B phosphorylation promotes localization of CNOT3 to the nucleus also suggests that the interaction between phosphorylated CNOT3 and ERK could be involved in stabilising phospho-ERK in the cytoplasmic and nuclear compartments and potentially in the transport of phospho-ERK into the nucleus.

**Figure 8.**
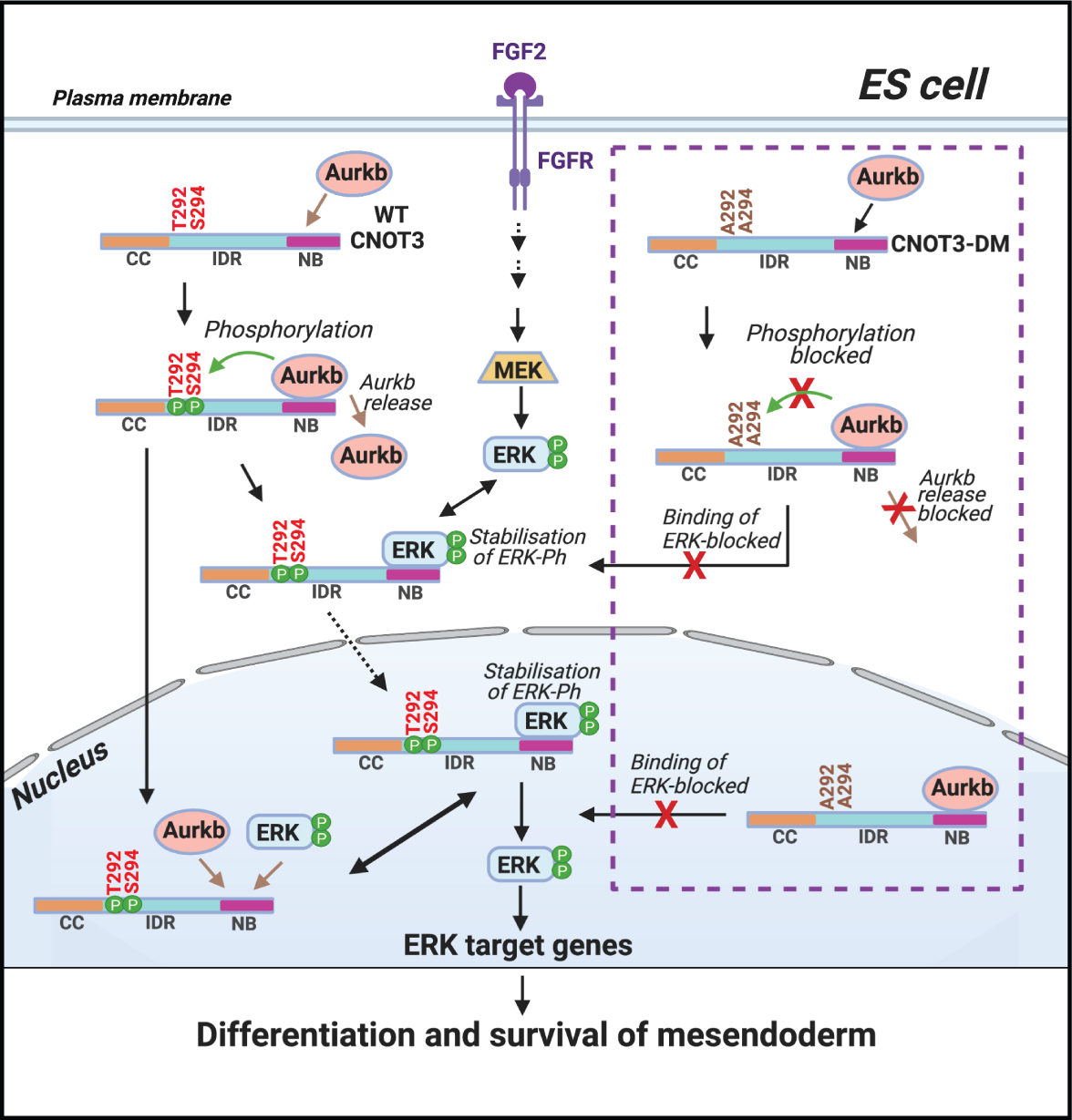
Schematic model showing the proposed regulation of ERK activity by Aurora B and CNOT3. Aurora B (AURKB) binds to the NOT box (NB) of CNOT3 (for clarity the additional synergistic contact with the amino-terminal coiled-coil (CC) domain is not shown). The bound Aurora B phosphorylates CNOT3-T292/S294, located adjacent to a nuclear localization signal in the intrinsically disordered region (IDR) of CNOT3 and promotes uptake of CNOT3 into the nucleus. Phosphorylation of CNOT3 also causes bound Aurora B to be released, allowing phospho-ERK (ERK-Ph) to bind to the NOT box. This results in stabilisation of ERK-Ph in the cytoplasm and nucleus and may also facilitate transport of ERK-Ph into the nucleus. Mutation of CNOT3-T292/S294 to A (CNOT3-DM) prevents phosphorylation of these residues by Aurora B (depicted inside the dashed box). This blocks release of Aurora B and prevents binding of ERK to the NOT box, resulting in destabilisation and downregulation of ERK-Ph.

In addition to these conclusions, the increased nuclear localisation of CNOT3 that we observe in an NSCLC cell line that was isolated from a lymph node metastasis opens up the possibility that Aurora B-dependent nuclear localisation of CNOT3 promotes EMT/MET in cancer cells, acting as a driver for metastasis. The known association of elevated Aurora B levels with lymph node metastasis in a number of cancers ^28–30^ and the well-established involvement of MAPK/ERK pathways in tumourigenesis and metastasis ^54–56^ suggests that the interaction of CNOT3, with Aurora B and ERK has important roles in the development and progression of human cancers.

Overall, our results provide evidence of a signaling axis between ERK and Aurora B with CNOT3 acting as an adaptor that links the the two kinases. Our findings suggest that Aurora B acts through CNOT3 to increase the level of phospho-ERK, thereby promoting survival and differentiation of ME cells. These observations indicate that the synergy between the three proteins plays an important role in regulating EMT during ME differentiation and gastrulation of the mouse embryo. The very strong reduction in survival and proliferation that we observe when the *Cnot3*-DM cells are induced to differentiate into ME suggests that Aurora B links up with the MAPK/ERK and Wnt pathways via CNOT3 to promote the explosive expansion of cell numbers that occurs in the embryo at the time of implantation and gastrulation.

## MATERIALS AND METHODS

### Cell Culture

E14 ESCs (female, 129/Ola) were obtained from the LMS Transgenics and ES cell facility. The ESCs were maintained on gelatin coated plates, at 37°C, 5% CO_2_, in KnockOut Dulbecco’s Modified Eagle’s Medium supplemented with 15% fetal bovine serum, 1x NEAA, 2 mM L-glutamine, 100 units/ml penicillin, 100 μg/ml streptomycin, 100 μM *β*ME (all reagents from Thermo Fisher Scientific, USA) and 1000 units/ml LIF (Merck, USA). HEK 293 cells and A549 cells (ATCC, USA) were maintained in Dulbecco’s Modified Eagle’s Medium supplemented with 10% fetal bovine serum, 2 mM L-glutamine, 100 units/ml penicillin and 100 μg/ml streptomycin. H1299 cells (ATCC, USA) were maintained in RPMI-1640 media supplemented with the same constituents as mentioned above.

### CNOT3 gene editing using CRISPR/Cas9

The guide RNA (gRNA) used to generate the *Cnot3*-T292A/S294A mutant ESCs was 5’-GATTTAGACTTGGACCCACC-3’. The gRNA was cloned into pX330 (Addgene, USA; plasmid: 42230) using forward primer 5’-CACCGGATTTAGACTTGGACCCACC-3’ and reverse primer 5’-AAACGGTGGGTCCAAGTCTAAATCC-3’ The 110 base paired single stranded donor DNA used to target *Cnot3* in exon 10 carrying the mutations was: 5’-ACTCTGAAGATGATAAGAAGAGAGGCCGATCTGCGGATGCTGAAGTCAGCCAGGTGGGTCCAAGTCTAAATCTGATGGTTTGTAACTTGTTTATTGCGTGGTCTCCAAAG-3’ Mouse ESCs (4×10^6^ cells) were transfected with 3 μg of pX330 plasmid carrying the gRNA, 4 μg of the donor ssDNA and 3 μg of a puromycin resistance plasmid (pCAG-puro^R^) using the Mouse ES Cell Nucleofector™ Kit (Lonza, Switzerland) following the manufacturer’s protocol. One day post transfection cells were subjected to puromycin selection (1.5 μg/ml) for 24 hours. A week after transfection, individual clones were picked and genotyped by allele specific nested PCR. Mutant genotypes were confirmed by sequencing.

### Embryoid bodies

Half a million cells (per well of a six well plate) were seeded on ultra-low attachment plates (Corning Costar, USA) and maintained in KnockOut Dulbecco’s Modified Eagle’s Medium supplemented with 15% fetal bovine serum, 1x NEAA, 2 mM L-glutamine, 100 units/ml penicillin, 100 μg/ml streptomycin, 100 μM *β*ME and were grown up to 14 days. Embryoid bodies (EBs) were fixed in 4% paraformaldehyde for twenty minutes followed by permeabilization with 0.5% Triton-X for twenty minutes.

Subsequently, EBs were blocked in 3% BSA for one hour and then incubated with primary antibodies overnight at 4°C. On the following day, the EBs were washed 3 times with PBS+0.1 % Tween-20 (PBST) and incubated for 1hour at room temperature with the appropriate secondary antibodies. Embryoid bodies were then washed three times in PBST and incubated with 1μg/ml DAPI (Merck, USA) in PBS for 45 minutes. Images were acquired using a SP5 confocal microscope with LAS X software (Leica Microsystems, Germany) and were analysed with Fiji ImageJ software (NIH, USA).

### Differentiation experiments

#### ESC differentiation into mesendoderm

Cells were plated at a density of 10000 cells/cm^2^ on gelatin-coated plates and incubated in DMEM/F12 KnockOut media containing 64 μg/ml of L Ascorbic acid-2-phosphate magnesium, 543 μg/ml of sodium bicarbonate, 1 μg/ml of heparin, 1x insulin-transferrin-selenium and 2mM Glutamine. For the differentiations, different combinations of signalling factors were added to the medium (see “Treatments of the cells with ligands and inhibitors” below). For time-lapse imaging of differentiation from day 3 to day 7, the plate was transferred to a Axiovert 200 microscope (Zeiss, Germany) with environmental chamber (Solent Scientific Ltd., UK) and motorised stage (ASI, USA) and images were collected at an interval of thirty minutes. Phase contrast images were acquired in DMIRE2 microscope (Leica Microsystems, Germany) using the MetaMorph software.

#### ESC differentiation to endoderm

Cells were plated at a density of 10000 cells/cm^2^ on gelatin-coated plates and incubated in high glucose DMEM with 15% FBS, 100 units/ml penicillin, 100 μg/ml streptomycin, 0.1 mM non-essential amino acids, 1mM MTG, 1x GlutaMAX and supplemented with 25 ng/ml of FGF2 (Merck, USA) and 10 μM of retinoic acid (Merck, USA) for 3 days ^57^.

#### ESC differentiation to mesoderm

Cells were plated at a density of 15000 cells/cm^2^ in gelatin + fibronectin-coated plates and incubated for 4 days in DMEM/F12-Neurobasal (1:1), N2, B27, 1x GlutaMAX, 100 units/ml penicillin, 100 μg/ml streptomycin, 0.1% *β*-ME and 30 ng/ml of Activin A (R&D Systems, USA).

#### ESC differentiation to ectoderm

Cells were plated at a density of 15000 cells/cm^2^ on gelatin-coated plates and incubated for 4 days in DMEM/F12-Neurobasal medium (1:1), 1x GlutaMAX, 100 units/ml penicillin, 100 μg/ml streptomycin, 100 μM *β*ME, B27 minus Vitamin A and N2.

### Treatment of cells with ligands and inhibitors

Aurora B inhibitor AZD1152 (Merck, USA)-200 nM; human recombinant Wnt3 (Cloud-Clone Corp, USA)-200 ng/ml; Activin A (R&D Systems, USA)-100 ng/ml; BMP4 (Merck, USA)-10 ng/ml; FGF2 (Merck, USA)-25ng/ml; CHIR99021(Merck, USA)-3μM, PD0325901 (Merck, USA)-500nM; U0126(Merck, USA)-10μM; AZD4547 (Abcam, UK)-5nM.

### Cell survival assay and Annexin V staining

The Annexin V staining was performed using the FITC Annexin V apoptosis detection kit with PI (BioLegend, USA). The cells were washed with PBS followed by washing with staining buffer and resuspension in binding buffer containing anti-Annexin V and PI and incubation for fifteen minutes at room temperature in the dark. The cells were finally washed with binding buffer and incubated with 1μg/ml DAPI in PBS for 5 minutes. Images were acquired using IX70 microscope (Olympus, Japan) with Micro-manager software.

Cells were stained with 0.2% crystal violet for a gross estimation of efficiency of differentiation. At different time points of differentiation live and dead cells were distinguished by trypan blue staining and counted manually. Cell survival was measured using Cell Proliferation Reagent WST-1 (Merck, USA) at a final dilution of 1:10 followed by incubation for half an hour and quantitation with a scanning multi-well spectrophotometer (SpectraMax-Molecular Devices, USA).

### Immunocytochemistry

All differentiated cells were grown in gelatin coated μ-slides (ibidi, Germany) for immunofluorescence. Cells were fixed in 4% paraformaldehyde for ten minutes, permeabilized with 0.5% Triton-X for fifteen minutes and blocked in 3% BSA for one hour. Following incubation with primary antibodies at 4°C overnight, the cells were washed with PBST and incubated for 1hour at room temperature with the appropriate secondary antibodies. DAPI was used to stain the nuclei. Images were acquired using a SP5 confocal microscope with LAS X software (Leica Microsystems, Germany) or IX70 microscope with Micromanager software.

Images were analysed with Fiji ImageJ software (NIH, USA). CellProfiler software (Broad Institute, USA) was used for the analysis of images from H1299 and A549 cancer cells. The antibodies used are listed in Supplementary Table 2.

### Expression plasmids

Full length Cnot3 cDNA (OriGene, USA) was cloned in pCDNA 3.2/V5/GW/D-TOPO by PCR addition of restriction sites SmaI/NotI following the manufacturer’s instructions. Single Cnot3-T292A, Cnot3-S294A mutant constructs and the T292A/S294A double mutant were generated by site directed mutagenesis and cloned as above. An 8 μg aliquot of each DNA construct was transfected into HEK 293 cells by Calcium Phosphate method. Deletion fragments Cnot3 *Δ*1-200, Cnot3 *Δ*651-700, Cnot3 *Δ*701-751, Cnot3 *Δ*651-751 were synthesized (Genewiz, UK) and cloned in pCDNA 3.2/V5/GW/D-TOPO using restriction sites KpnI/AscI. pCDNA3-T7-ERK1 (Addgene, USA; plasmid:14440) and full-length Aurora B cDNA (Dharmacon-Horizon Discovery, UK) was cloned in pGEX-4T1.

### Immunoprecipitation (IP)

Cells were harvested in IP Lysis Buffer (50 mM Tris-HCl, pH 7.4, 150 mM NaCl, 10% glycerol, 1% Nonidet P-40, 1.0 mM EDTA) with Complete™ protease inhibitor cocktail (Merck, USA). In all cases, 500 μg of total protein was used. Extracted proteins were immunoprecipitated with Protein A Sepharose CL-4B beads (GE Healthcare, USA) and desired primary antibodies. The immunocomplexes were eluted by boiling with 2x SDS loading buffer.

### Protein extractions from cells

To obtain the whole cell extracts, cells were washed with ice cold PBS and the cell pellet was reuspended in Tris lysis buffer (50 mM Tris/HCl pH 7.5, 150 mM NaCl, 1% Triton X-100, 1 mM DTT, 1 mM Na_3_VO_4_) with Complete™ protease inhibitor cocktail (Merck, USA).

For preparation of cytoplasmic and nuclear extracts, cells were harvested and re-suspended in harvest buffer containing 10 mM HEPES (pH 7.9), 50 mM NaCl, 0.5 M Sucrose, 0.1 mM EDTA, 0.5% Triton-X 100 and with Complete™ protease inhibitor cocktail (Merck). After obtaining the cytoplasmic extract, the nuclear pellet was further washed with wash buffer/Buffer A (10 mM HEPES (pH 7.9), 10 mM KCL, 0.1 mM EDTA and 0.1 mM EGTA), then re-suspended in Buffer C (10 mM HEPES (pH 7.9), 500 mM NaCl, 0.1 mM EDTA, 0.1 mM EGTA, 0.1% NP40 and protease inhibitor cocktail to extract the nuclear proteins.

### Immunoblotting

Immunoprecipitated proteins in loading buffer or equal amounts of proteins obtained from cell extracts (diluted in 5x SDS loading buffer) were boiled for 5 min and subjected to SDS PAGE. Gels were transferred to nitrocellulose membranes (GE Healthcare, USA). Membranes were blocked with 5% bovine serum albumin (BSA) or 5% milk for 1hour at room temperature and incubated with the desired primary antibodies overnight at 4°C. On the following day, membranes were washed 3 times with TBS-Tween 20, incubated with the appropriate secondary antibodies (dilution 1:10000) for 1hour at room temperature, washed and developed using Crescendo ECL (Merck, USA) using X ray films on Photon Imaging System (UK) or Amersham Imager 680 (GE healthcare, USA).

### Protein purification in bacteria

pGEX-4T1-Aurora B and pGEX-4T1-ERK1 were transformed into competent BL21 E.coli cells. To induce protein expression, β-D-1 thiogalactopyranoside (IPTG) (Merck, USA) was added at a concentration of 0.5 mM and the cells were grown for 3 hours at 37°C. The cell pellets were resuspended in PBS+0.5% Triton-X 100 supplemented with Complete™ protease inhibitor cocktail (Merck, USA) and frozen overnight at −80 °C for lysis. The following day, the mix was incubated with Lysozome (1mg/ml) followed by sonication at intervals of 15 seconds on and 45 seconds off for 5 minutes. The clear supernatants were collected and Sepharose High performance (GSH) beads (GE Healthcare, USA) was added followed by overnight incubation at 4°C. Beads were then washed with PBS+0.5% Triton-X 100 and the GST fusion proteins were eluted with elution buffer (10mM glutathione reduced, 5% glycerol in 50 mM Tris-Cl pH 8.0) for 15 minutes at room temperature followed by dialysis in 50 mM Tris-Cl pH 8.0, 0.15 M NaCl, 10% glycerol, 0.5 mM DTT and 0.5mM PMSF.

### GST pulldown assay

In order to perform the GST pulldown with GST-Aurora B and GST-ERK1 with Cnot3 and the deletion mutants of Cnot3, 0.5 μg of pCDNA-*Cnot3*, pCDNA-Cnot3 *Δ*1-200, pCDNA-Cnot3 *Δ*651-700, pCDNA-Cnot3 *Δ*701-751 and pCDNA-Cnot3*Δ*651-751 were *in vitro* transcribed/translated using a TNT^®^ Quick Coupled Transcription/Translation kit (Promega, USA) according to manufacturer’s instructions using 10μCi of ^35^S-Methionine to radiolabel the proteins. 1μg of GST or GST-Aurora B or GST-ERK was added to GSH beads (GE Healthcare, USA) in binding buffer (50 mM Tris-cl pH 8.0, 150 mM monopotassium glutamate, 1mM EDTA, 0.1 % Igepal CAL630, 5% glycerol, 0.2% BSA) and supplemented with Complete™ protease inhibitor cocktail (Merck, USA), and incubated for two hours at 4°C. The beads were then washed and 5 μl of the *in vitro* transcribed/translated CNOT3 was incubated with the beads overnight at 4°C. The beads were then washed with the binding buffer and the proteins were eluted by boiling in loading buffer. The eluted proteins were subjected to SDS-PAGE, stained, dried for one hour at 80°C and exposed overnight to Phosphor screen in a cassette (GE healthcare, USA). Images were captured in Fujifilm FLA 5100 scanner (Japan) using the Fujifilm FLA-5000 software.

### *In vitro* kinase assay

pCDNA-Cnot3-T292A, pCDNA-Cnot3-S294A, pCDNA-T292A/S294A and pCDNA-Cnot3 *Δ*651-751 constructs were *in vitro* transcribed/translated as described above and the resultant V5 tagged proteins were purified using V5-tagged protein purification kit ver.2 (MBL International, USA). Purified proteins were used as substrate and incubated with 80 ng of GST-Aurora B (ProQuinase, Germany) for 30 min at 30°C in phosphorylation buffer (50 mM TrisHCl pH 7.5, 10 mM MgCl_2_, 500 μM ATP, 1 mM DTT, 5 mM NaF, 1 μl of ^32^P*γ*-ATP 10μCi). Reactions were stopped by addition of SDS loading buffer, boiled for 5 min and subsequently run on SDS PAGE. The gel was dried and exposed to X ray film.

### Flow cytometry

Cells were trypsinised, washed twice with PBS, fixed with 70% ethanol for 30 min on ice and washed twice with 2% FCS-PBS. Subsequently, cells were resuspended in PBS containing 1μg/ml RNAse A (Thermo Fisher Scientific, USA), 50 μg/ml propidium iodide and 0.05% NP40, incubated 20 min at room temperature in the dark followed by analysis. Anaysis was performed on an LSRII Flow Cytometer (Becton-Dickinson, USA) and data was analysed using FlowJo Software (Becton-Dickinson, USA).

### RNA sequencing

Total RNA was extracted from 2×10^6^ cells (3 biological replicates for wild-type and CNOT3-DM cells) using Trizol reagent (Thermo Fisher Scientific, USA) following the manufacturer’s instructions. A 2 μl aliquot of a 1:10 dilution of the ERCC RNA Spike-in Mix (Thermo Fisher Scientific) were added to each sample. The quality of the extracted RNA was analysed using the RNA 6000 Nano kit on a 2100 Bioanalyzer (Agilent, USA). An aliquot of 500 ng of RNA was used to prepare a polyadenylated RNA library with the TruSeq Stranded mRNA Library Prep Kit (Illumina, USA) following the manufacturer’s protocol. RNA libraries were sequenced in one lane by multiplexing on an Illumina HiSeq 2500 sequencer with a 100 bp read output. RNAseq reads were aligned against to Ensembl Mouse genome (NCBIM37) reference sequence assembly and transcript annotation that obtained from Illumina iGenomes (https://support.illumina.com/sequencing/sequencing_software/igenome.html) and to ERCC reference with Tophat2 (2.0.11) ^58^. Gene-based read counts and ERCC counts were obtained using featureCounts function from Rsubread Bioconductor package ^59, 60^. Differentially expressed gene analysis was performed with DESeq2 Bioconductor package ^61^ after normalising the data against ERCC with RUVseq Bioconductor package ^62^. Differentially expressed genes were defined with Benjamini-Hochberg adjusted p-value<0.05 and fold change ratio > 1.5. Gene ontology analysis was performed with goseq Bioconductor package ^63^. After converting mouse gene symbol to human gene symbol using the report of Human and Mouse Homology retrieved from Mouse Genome Informatics (MGI, www.informatics.jax.org), Gene Set Enrichment Analysis (GSEA) ^64, 65^ was then performed with GseaPreranked tool using Hallmarks gene set (h.all.v5.2.symbols.gmt).

### Proximity ligation assay (PLA)

PLA was performed using the Duolink In Situ Red Starter Kit Mouse/Rabbit (Merck, USA) following manufacturer’s instructions. Images were acquired using a SP5 confocal microscope with LAS X software (Leica microsystems, Germany). PLA dots were analysed and quantified using Imaris-Bitplane software (Oxford Instruments, UK). Three-dimensional segmentation and digital image reconstructions of the cells were carried out using Imaris Spots and Surfaces packages.

### Mass spectrometry

Peptides were separated using an Ultimate 3000 RSLC nano liquid chromatography system (Thermo Fisher Scientific, USA) coupled to an LTQ Velos Orbitrap mass spectrometer (Thermo Fisher Scientific, USA) via a Dionex nano-esi source. Eluted peptides were analysed by the LTQ Velos operating in positive polarity using a data-dependent acquisition mode. Ions for fragmentation were determined from an initial MS1 survey scan at 60000 resolution (at m/z 200).

Ion Trap CID (collisional induced dissociation), Ion Trap CID-MSA (MultiStage Activation) and HCD (Higher energy collisional induced dissociation) were carried out concurrently on the top 3 most abundant ions. For CID fragmentation methods MS1 and MS2 MSn AGC targets set to 1e6 and 1e4 for 1 microscan and a maximum injection time of 500ms and 100ms respectively. A survey scan m/z range of 350 – 1500 was used, with a normalised collision energy set to 35% for both CID and HCD, charge state rejection enabled for +1 ions and a minimum threshold for triggering fragmentation of 500 counts.

### RNA isolation and quantitative RT-PCR (qPCR)

Total RNA was isolated with TRIzol reagent (Thermo Fisher Scientific, USA) following manufacturer’s instructions. cDNA was synthesized with RevertAid First Strand cDNA Synthesis Kit (Thermo Fisher Scientific, USA) using 200 ng of RNA following the manufacturer’s protocol. qPCRs were performed using Sensimix SYBR NORox SYBR GREEN (Bioline, UK). Each PCR was performed in duplicate using 1 μl of cDNA (from 20 μl reaction) and 200 nM primer concentration. Gene expression was determined relative to *MLN 51* transcript levels. The primers used in this analysis are listed in Supplementary Table 1.

### Statistical analysis

All statistical analyses were performed with GraphPad Prism software (GraphPad, USA). The statistical tests used in each experiment and significances are indicated in the corresponding figure legends.

## Data availability

The RNA-seq data have been deposited at GEO under accession number GSE138213. Reviewer token for accessing the data: gtynmayutbiphqf.

## ACKNOWLEDGEMENTS

We thank Nicola Festuccia, Aida di Gregorio and Sheila Xie for advice and helpful discussions, Holger Kramer and Alex Montoya for assistance with mass spectrometry data analysis and Lepakshi Ranjha for advice on the ^35^S pull-down and in vitro kinase assays. We also thank the staff of the LMS Genomics Facility for HT sequencing, and the LMS/NIHR Imperial Biomedical Research Centre Flow Cytometry Facility for support.

## Funding

This research was funded by the Medical Research Council UK.

## Conflict of interest

The authors declare that they have no conflict of interest.

## Legend to Supplementary Video 1

Time-lapse imaging of BMP4/FGF2-induced differentiation of wild-type and *Cnot3*-DM ESCs. Images were collected at 30 minute intervals from day 3 to day 7 of the differentiation (see Methods). Display rate: 15 frames per second. Scale-bar = 100 μM.

## Notes

### Competing Interest Statement

The authors have declared no competing interest.

