## Supplementary information for "A signalling axis involving CNOT3, Aurora B and ERK promotes differentiation and survival of mesendodermal progenitor cells"

**Sarkar et al.**

### SUPPLEMENTAL INFORMATION

#### Figure S1

**A**

```
10         20         30         40         50
MADKRKLQGE IDRCLKKVSE GVEQFEDIWQ KLHNAANANQ KEKYEADLKK
        60         70         80         90        100
EIKKLQRLRD QIKTWVASNE IKDKRQLIEN RKLIETQMER FKVVERETKT
       110        120        130        140        150
KAYSKEGLGL AQKVDPAQKE KEEVGQWLTN TIDTLNMQVD QFESEVESLS
       160        170        180        190        200
VQTRKKKGDK DKQDRIEGLK RHIEKHRYHV RMLETILRML DNDSILVDAI
       210        220        230        240        250
RKIKDDVEYY VDSSQDPDFE ENEFLYDDLD LEDIPQALVA TSPPSHSHME
       260        270        280        290   *  * 300
DEIFNQSSST PTSTTSSSPI PPSPANCTTE NSEDDKKRGR STDSEVSQSP
       310        320        330        340        350
AKNGSKPVHS NQHPQSPAVP PTYPSGPPPT TSALSSTPGN NGASTPAAPT
       360        370        380        390        400
SALGPKASPA PSHNSGTPAP YAQAVAPPNA SGPSNAQPRP PSAQPSGGSG
       410        420        430        440        450
GGSGGSSSNS NSGTGGGAGK QNGATSYSSV VADSPAEVTL SSSGGSSASS
       460        470        480        490        500
QALGPTSGPH NPAPSTSKES STAAPSGAGN VASGSGNNSG GPSLLVPLPV
       510        520        530        540        550
NPPSSPTPSF SEAKAAGTLL NGPPQFSTTP EIKAPEPLSS LKSMAERAAI
       560        570        580        590        600
SSGIEDPVPT LHLTDRDIIL SSTSAPPTSS QPPLQLSEVN IPLSLGVCPL
       610        620        630        640        650
GPVSLTKEQL YQQAMEEAAW HHMPHPSDSE RIRQYLPRNP CPTPPYHHQM
       660        670        680        690        700
PPPHSDTVEF YQRLSTETLF FIFYYLEGTK AQYLAAKALK KQSWRFHTKY
       710        720        730        740        750
MMWFQRHEEP KTITDEFEQG TYIYFDYEKW GQRKKEGFTF EYRYLEDRDLQ
```

Predicted coiled-coil domain (1-241)

Intrinsically disordered region (242-605)

NAR/CS

NOT box (654-751)

**B**

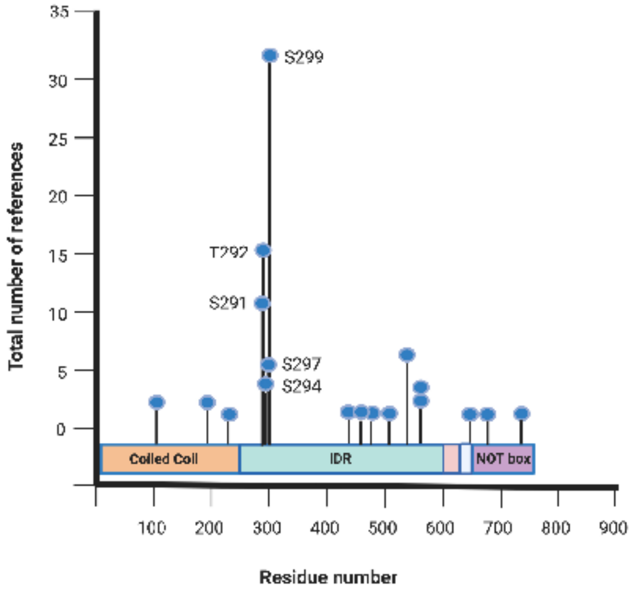

**Supplementary Figure 1: A**. Sequence of the mouse CNOT3 protein (UNIPROT Q8K0V4). Vertical blue lines delineate the domains of the protein that are shown in Figure 1B. The green-shaded box indicates the the NOT box region, which is required for the interaction between CNOT3 and Aurora B (Figure 1C). Aurora B consensus phosphorylation sites are underlined. The red box and red text indicate the putative nuclear localization sequence. * indicates residues 292 and 294, which were mutated in this study.
**B.** Phosphorylation sites detected on the CNOT3 protein by *in vivo* proteomic discovery mass pectrometry. Data was obtained from PhosphoSite® (Hornbeck et al., 2015). The data shown is for the human CNOT3 protein, which is 95% identical to mouse CNOT3. With the exception of one residue (E79), all of the residues that vary between human and mouse are located within the intrinsically disordered region between residues 329 and 604 of the mouse sequence. Blue circles indicate phosphorylated residues. The y-axis shows the number of records where the modification was assigned using proteomic discovery mass spectrometry. The numbered residues show a phosphorylation hotspot that extends from positions 291-299.

### Figure S2

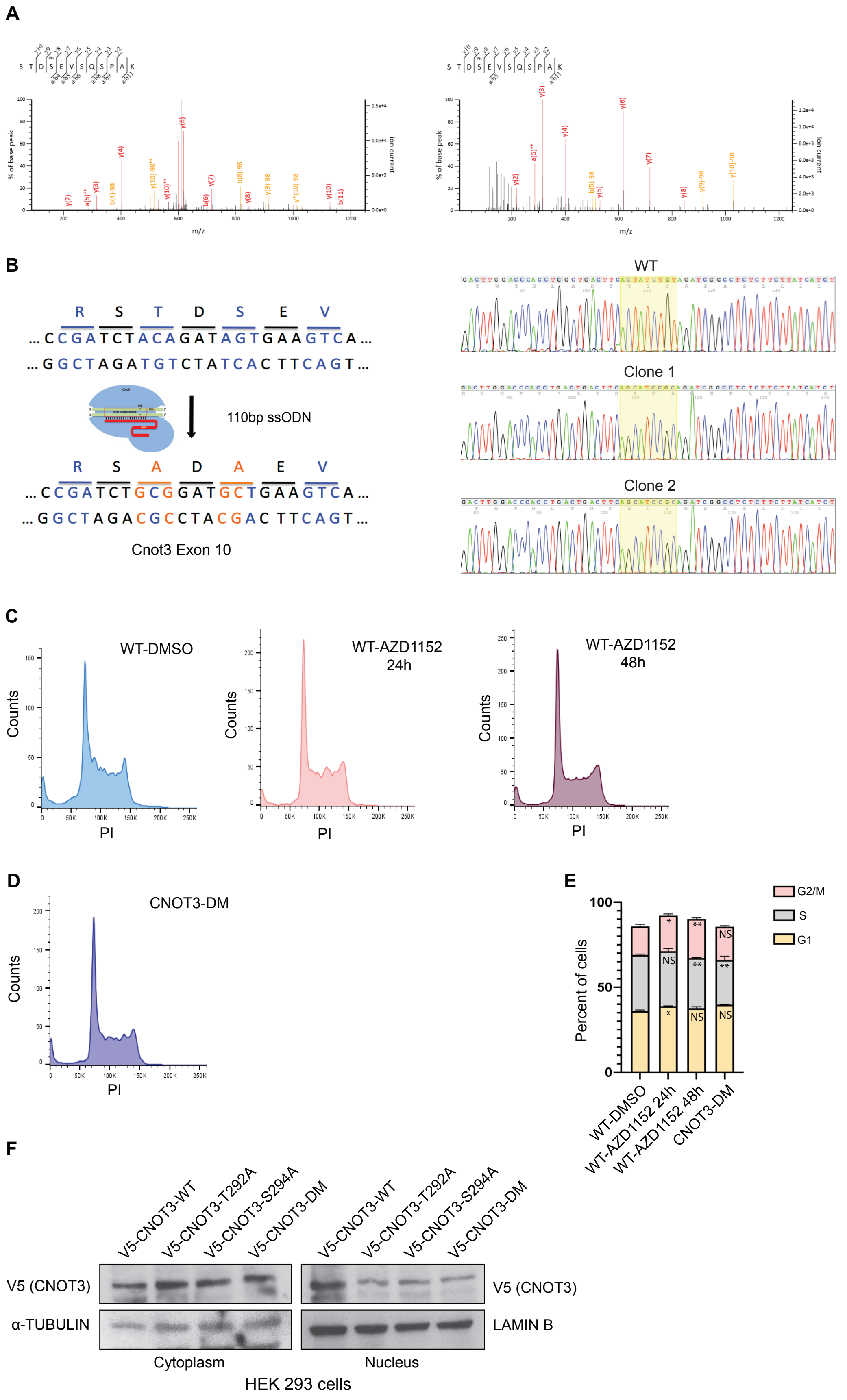
A
S T D S E V S Q S P A K
% of base peak
Ion current
m/z
B
R S T D S E V
... C C G A T C T A C A G A T A G T G A A G T C A ...
... G G C T A G A T G T C T A T C A C T T C A G T ...
110bp ssODN
R S A D A E V
... C C G A T C T G C G G A T G C T G A A G T C A ...
... G G C T A G A C G C C T A C G A C T T C A G T ...
Cnot3 Exon 10
WT
Clone 1
Clone 2
C
WT-DMSO
WT-AZD1152 24h
WT-AZD1152 48h
Counts
PI
D
CNOT3-DM
Counts
PI
E
Percent of cells
G2/M
S
G1
NS
WT-DMSO
WT-AZD1152 24h
WT-AZD1152 48h
CNOT3-DM
F
V5-CNOT3-WT
V5-CNOT3-T292A
V5-CNOT3-S294A
V5-CNOT3-DM
V5 (CNOT3)
α-TUBULIN
LAMIN B
Cytoplasm
Nucleus
HEK 293 cells

**Supplementary Figure 2: Identification of Aurora B phosphorylation sites in the CNOT3 protein**

**A.** MS/MS fragmentation spectrum generated by ion trap collision-induced dissociation (CID) of doubly charged precursor ion with m/z 658.27 identifying CNOT3 tryptic peptide 291-302 (STDSEVSQSPAK). The spectrum presented two fragments which were as follows: Left panel: query number 2609 with assigned b- and y-ions; MASCOT score: 54; expectation value: $1.8 \times 10^{-5}$; neutral loss of $H_3PO_4$ (-97.98 Da) from precursor ion detected at m/z 609.49; localisation probability for phosphorylation of Ser291: 3.0%, Thr292: 3.0%, Ser294 is 93.9%. Right panel: query number 2608 with assigned b- and y-ions; MASCOT score: 53; expectation value: $3.6 \times 10^{-5}$; localisation probability for phosphorylation of Ser291: 25.6%, Thr292: 25.6%, Ser294: 48.4%.

**B.** Left panel: Schematic representation showing use of CRISPR/Cas9 targeting of *Cnot3* in exon 10 to mutate Threonine 292 and Serine 294 to alanine, using the guide RNA described in Methods and a 110 bp single-stranded donor oligonucleotides (ssODN) carrying the mutations. Right panel: Sequence images showing the wild-type (WT) sequence and the mutated sequences obtained from two CRISPR generated clones.

**C.** FACS analysis of the cell cycle profiles of ESCs incubated for 24 and 48 hours with the Aurora B inhibitor AZD1152 or with vehicle (DMSO) and stained with propidium iodide (PI).

**D.** FACS analysis of the cell cycle profile of *Cnot3*-DM ESCs.

**E.** Histograms showing the percentage of cells measured at different phases of the cell cycle in (**c**) and (**d**). Mean ± SEM; P values (with respect to WT-DMSO) calculated by unpaired *t*-test, ns = non-significant, *n*=3.

**F.** Representative immunoblot analysis of V5-tagged CNOT3 carried out on cytoplasmic and nuclear extracts from HEK 293 cells transfected with WT *Cnot3*, *Cnot3*-T292A, *Cnot3*-S294A, or *Cnot3*-DM expression constructs. Cells were harvested 48 hours post-transfection. α-Tubulin and Lamin B were used as loading controls for the cytoplasm and nucleus respectively.

#### Figure S3

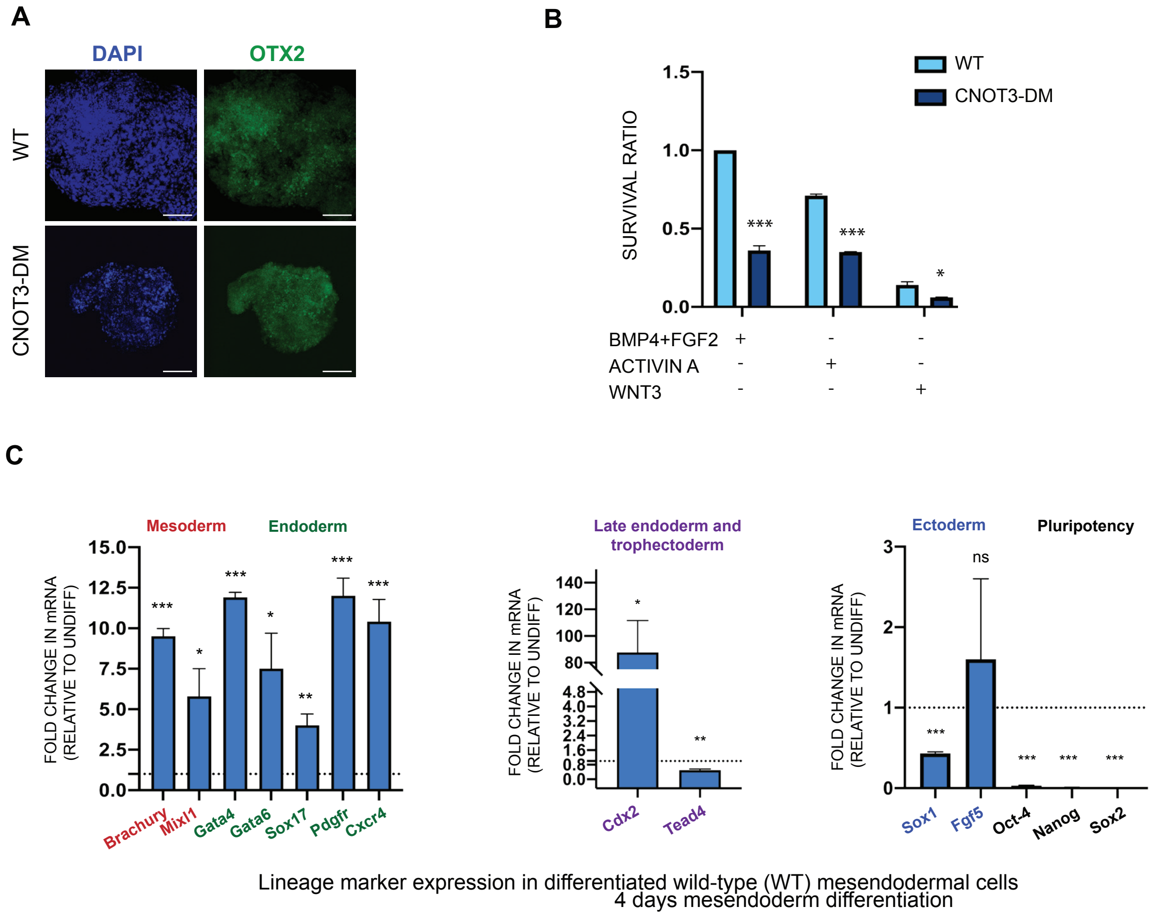


**Supplementary Figure 3: Differentiation and survival of *Cnot3*-DM mesendoderm**
**A.** Representative confocal images of 8-day embryoid bodies (EBs) derived from wild-type (WT) and *Cnot3*-DM ESCs and subjected to immunostaining for the ectodermal marker OTX2 (green). Nuclei were counterstained with DAPI; scale bar: 100 µm.
**B.** Survival of WT and *Cnot3*-DM ESCs over 4 days of differentiation induced in the presence of BMP4+FGF2 or Activin A or Wnt3 and the cell survival was determined using WST-1 reagent on day 4 of the differentiation. Histograms show survival ratios calculated relative to the values obtained for wild-type (WT) cells treated with BMP4+FGF2, which was assigned a value of 1. Mean ± SEM, significance calculated using unpaired t-test *P<0.05, ***P<0.001, n=3.
**C.** Analysis of mRNA levels of lineage marker genes after BMP4+FGF2 induced differentiation of WT ESCs for 4 days. Values for the undifferentiated (undiff) ESCs were set at 1 as indicated by a horizontal line in each histogram. *Cdx2* is a marker for late endoderm and trophectoderm. *Tead4* is a marker for early trophectoderm. Mean ± SEM; *n*=3. P values are calculated by unpaired *t*-test, *P<0.05, **P<0.01, ***P<0.001, ns = non significant.

#### Figure S4

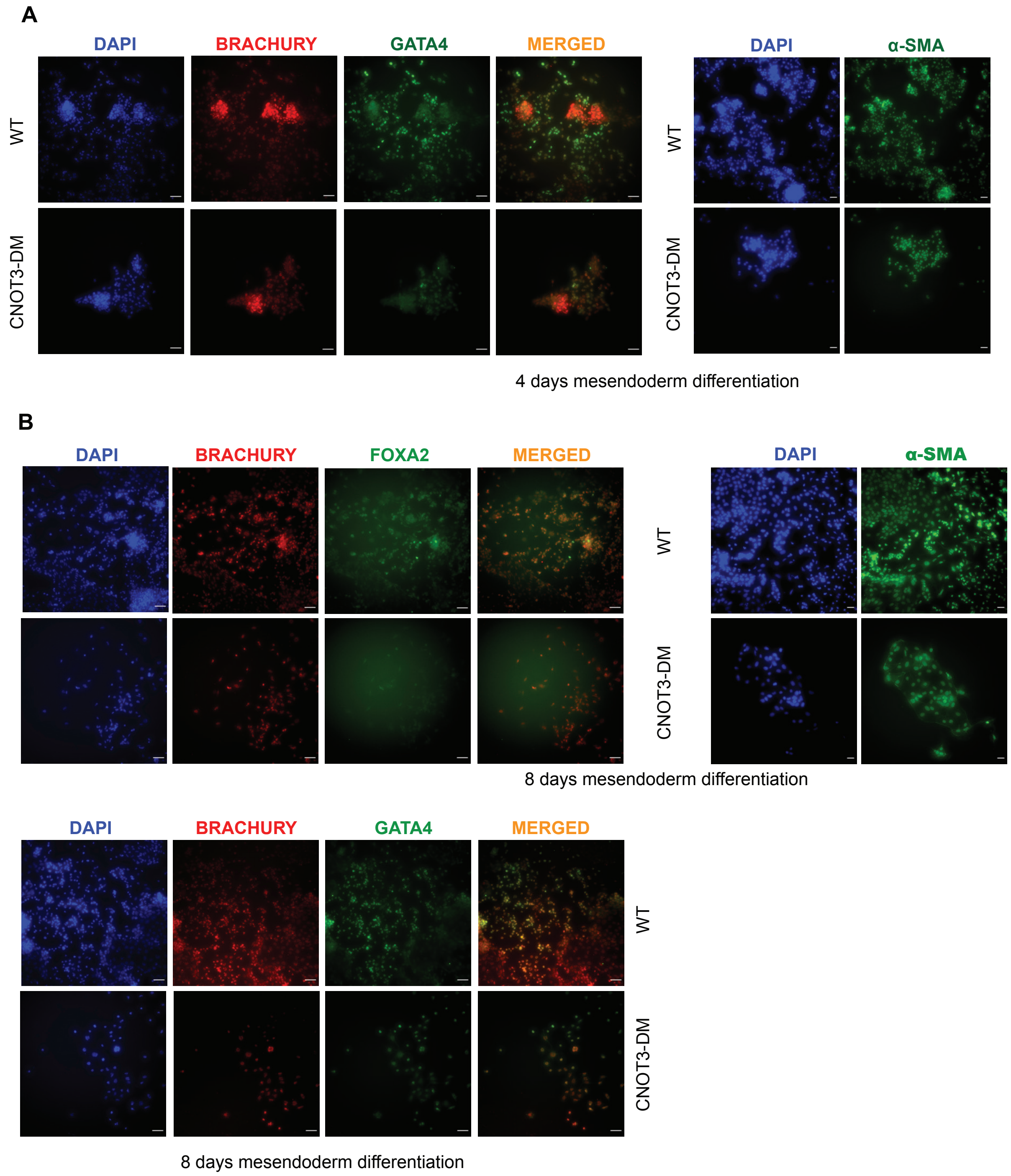


**Supplementary Figure 4: CNOT3-T292/S294 phosphorylation promotes survival of differentiating mesendodermal cells.**

**A.** Staining for ME markers shows that differentiation of *Cnot3*-DM ESCs with BMP4 + FGF2 for 4 days generates reduced numbers of ME cells. Left panel: Representative IF analyses of staining of differentiated cells for the mesodermal marker Brachury (*red*) and endodermal marker GATA4 (green) carried out on ME cells differentiated from WT and *Cnot3*-DM ESCs by incubating them with BMP4 + FGF2. Right panel: staining for the additional mesodermal marker smooth muscle actin (SMA) (green). Nuclei were stained with DAPI. Merged red and green images show the ME cells. Scale bars in (C) = 100 μm; scale bar in (D) = 60 μm. For IF analysis at 8 days of differentiation, see Figure S4A.

**B.** Representative immunofluorescence analysis images of ME cells generated by inducing differentiation of WT and *Cnot3*-DM ESCs with BMP4 + FGF2 for 8 days (left panel: scale bar: 100 μm). Top left panel: staining for the mesodermal marker Brachury (*red*) and endodermal marker FOXA2 (green). Bottom panel: staining for Brachyury (red) and the endodermal marker GATA4. Merged images show dual staining of the mesendodermal cells. Top right panel: staining for the mesodermal marker α-SMA (scale bar: 60 μm). Nuclei were counterstained with DAPI.

### Figure S5

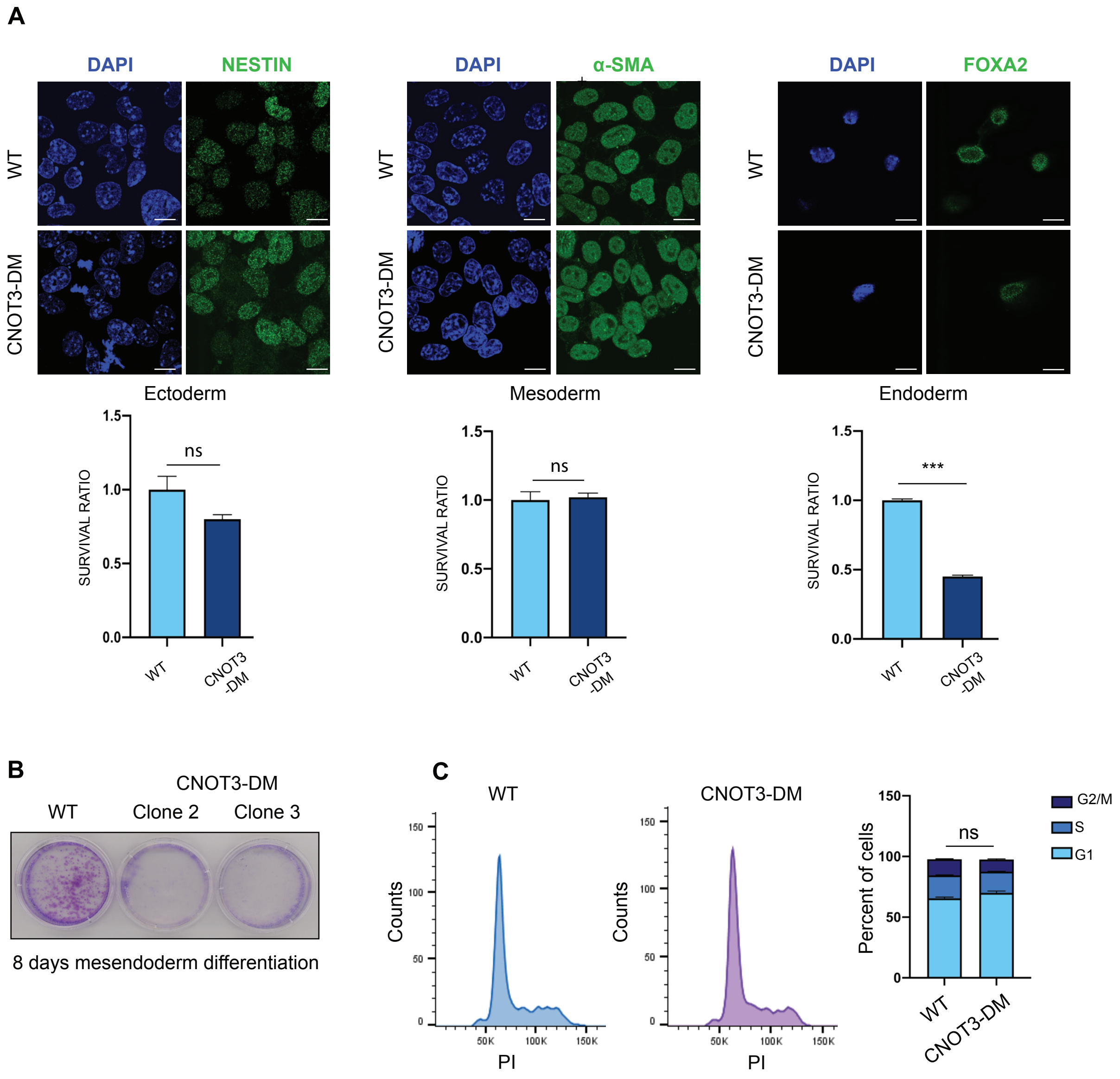


**Supplementary Figure 5: Differentiation of *Cnot3*-DM ESCs into the three germ layers.**

**A.** Top panels: Representative confocal images of differentiation of wild-type (WT) and *Cnot3*-DM ESCs into ectoderm (3 days; immunostained for Nestin (*green*)), Activin A induced mesoderm (3 days; immunostained for α-SMA (*green*)) and FGF2+retinoic acid induced endoderm (3 days; immunostained for FOXA2 (*green*). Nuclei were counterstained with DAPI; scale bar: 12 μm. (See Methods). Bottom panels: Graphs represent the ratio of suriving *Cnot3*-DM cells versus WT cells after 3 days of differentiation into each lineage. Cell survival was measured using WST-1 reagent (see Methods). Mean ± SEM; (P values calculated by unpaired t-test, ***P<0.001. ns = non-significant, n=3).

**B.** Crystal violet staining showing the efficiency of BMP4+FGF2 induced mesendoderm (ME) differentiation of wild-type (WT) ESCs and two additional clones of *Cnot3*-DM ESCs after 4 days of differentiation.

**C.** Cell cycle analysis of ME cells obtained by inducing differentiation of WT and *Cnot3*-DM ESCs with BMP4 + FGF2 for 4 days. Histograms represent the percentage of cells at different phases of the cell cycle. Mean ± SEM; P values calculated by unpaired *t*-test, ns = non-significant, *n*=3.

### Figure S6

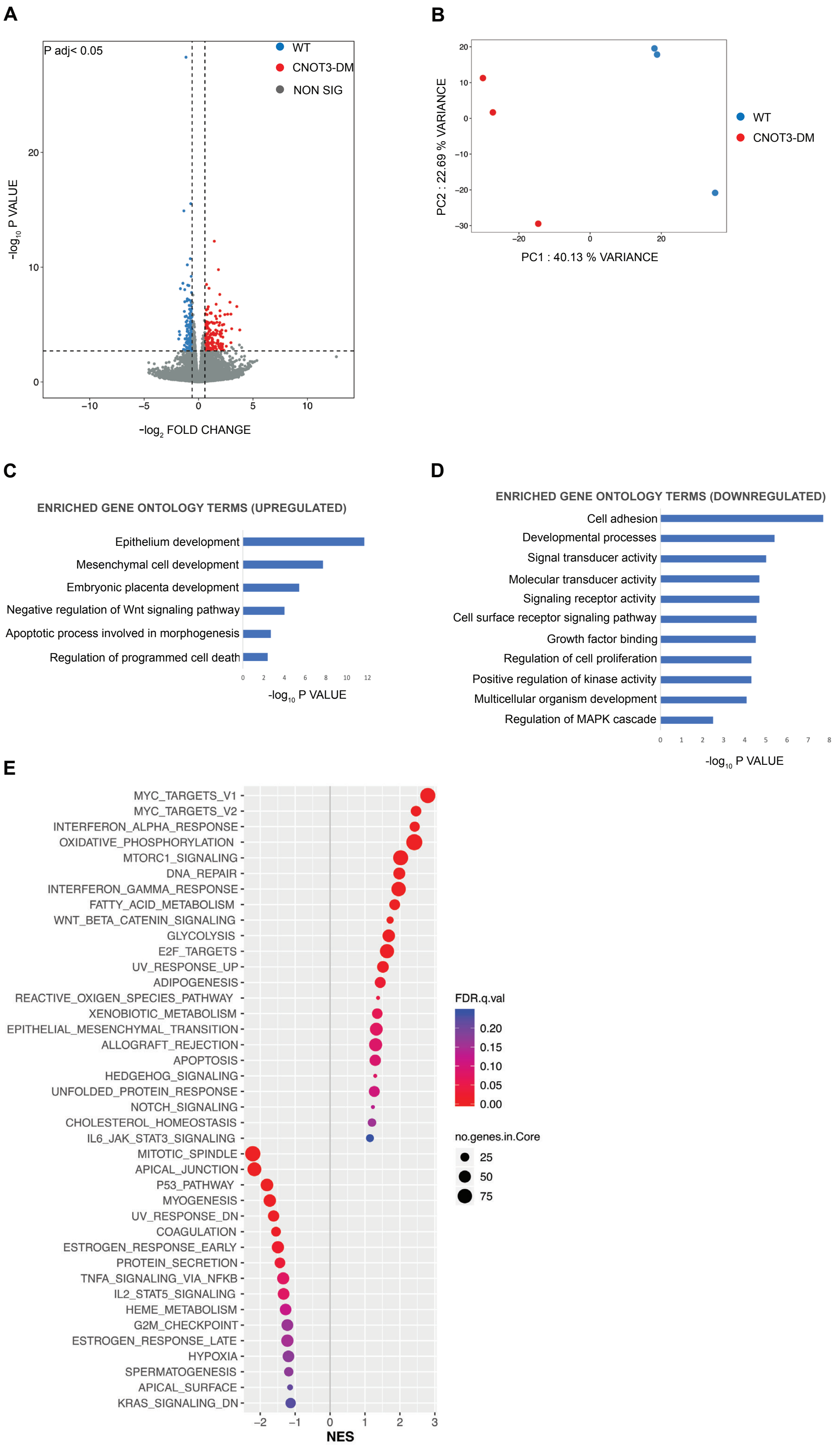
A
P adj< 0.05
WT
CNOT3-DM
NON SIG
-log10 P VALUE
-log2 FOLD CHANGE
B
PC2 : 22.69 % VARIANCE
PC1 : 40.13 % VARIANCE
WT
CNOT3-DM
C
ENRICHED GENE ONTOLOGY TERMS (UPREGULATED)
Epithelium development
Mesenchymal cell development
Embryonic placenta development
Negative regulation of Wnt signaling pathway
Apoptotic process involved in morphogenesis
Regulation of programmed cell death
-log10 P VALUE
D
ENRICHED GENE ONTOLOGY TERMS (DOWNREGULATED)
Cell adhesion
Developmental processes
Signal transducer activity
Molecular transducer activity
Signaling receptor activity
Cell surface receptor signaling pathway
Growth factor binding
Regulation of cell proliferation
Positive regulation of kinase activity
Multicellular organism development
Regulation of MAPK cascade
-log10 P VALUE
E
MYC_TARGETS_V1
MYC_TARGETS_V2
INTERFERON_ALPHA_RESPONSE
OXIDATIVE_PHOSPHORYLATION
MTORC1_SIGNALING
DNA_REPAIR
INTERFERON_GAMMA_RESPONSE
FATTY_ACID_METABOLISM
WNT_BETA_CATENIN_SIGNALING
GLYCOLYSIS
E2F_TARGETS
UV_RESPONSE_UP
ADIPOGENESIS
REACTIVE_OXIGEN_SPECIES_PATHWAY
XENOBIOTIC_METABOLISM
EPITHELIAL_MESENCHYMAL_TRANSITION
ALLOGRAFT_REJECTION
APOPTOSIS
HEDGEHOG_SIGNALING
UNFOLDED_PROTEIN_RESPONSE
NOTCH_SIGNALING
CHOLESTEROL_HOMEOSTASIS
IL6_JAK_STAT3_SIGNALING
MITOTIC_SPINDLE
APICAL_JUNCTION
P53_PATHWAY
MYOGENESIS
UV_RESPONSE_DN
COAGULATION
ESTROGEN_RESPONSE_EARLY
PROTEIN_SECRETION
TNFA_SIGNALING_VIA_NFKB
IL2_STAT5_SIGNALING
HEME_METABOLISM
G2M_CHECKPOINT
ESTROGEN_RESPONSE_LATE
HYPOXIA
SPERMATOGENESIS
APICAL_SURFACE
KRAS_SIGNALING_DN
NES
FDR.q.val
0.20
0.15
0.10
0.05
0.00
no.genes.in.Core
25
50
75

**Supplementary Figure 6: Transcriptome profiling by RNA sequencing in differentiating wild-type and *Cnot3*-DM mesendoderm cells**.

**A. and B.** RNA-seq analysis was used to compare gene expression patterns of ME cells obtained by differentiating wild-type (WT) and *Cnot3*-DM ESCs for 4 days with BMP4 + FGF2.

**A.** Volcano plot of RNA-Seq transcriptome data showing differentially expressed genes after 4 days of BMP4 + FGF2 induced mesendoderm differentiation of WT and *Cnot3*-DM ESCs. Fold change ratio >1.5; adjusted P value <0.05. Three biological replicates for each were analyzed.

**B.** Principal component plot of the RNA-seq data showing PC1 and PC2 and the percent variance obtained for each. This reveals separate clustering of the genes that are differentially expressed between the WT and *Cnot3*-DM-derived 4-day differentiated mesendodermal cells.

**C.** and **D.** Selected gene ontology (GO) profiles obtained out using the GOSeq Bioconductor package and showing upregulated genes (C) and downregulated genes (D), which are involved in signalling pathways that have key roles in developmental processes. Adjusted P value <0.05.

**E.** GSEA analysis showing Hallmark gene sets that were significantly altered in the differentiated cells. See also Supplementary Tables S3A and S3B for the differentially regulated genes. The GO and GSEA analysis both showed that expression of genes involved in cell death and apoptosis was upregulated (C and E). The negative regulator of Wnt signalling *Axin2*, and *TGFβ*, which is part of the apoptosis program, were also elevated. Additionally, the GO analysis revealed downregulation of gene expression programs that are involved in cell adhesion and developmental processes (D). The changes included downregulation of signalling molecules such as Wnt6. Bcl2, a key cell survival factor downstream of ERK signalling was also downregulated.

### Figure S7

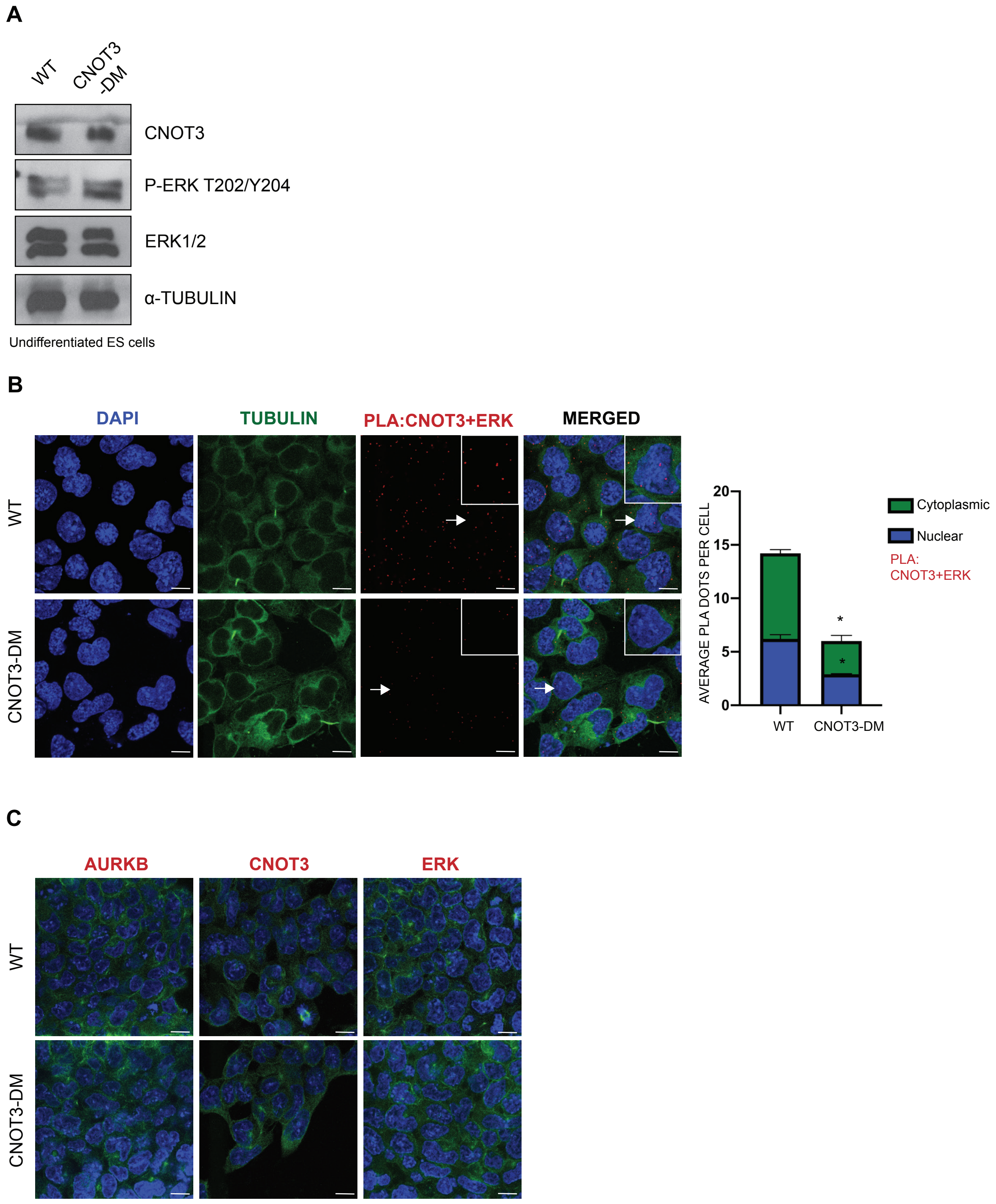
A
WT
CNOT3 -DM
CNOT3
P-ERK T202/Y204
ERK1/2
α-TUBULIN
Undifferentiated ES cells
B
DAPI
TUBULIN
PLA:CNOT3+ERK
MERGED
WT
CNOT3-DM
AVERAGE PLA DOTS PER CELL
20
15
10
5
0
WT
CNOT3-DM
Cytoplasmic
Nuclear
PLA: CNOT3+ERK
*
*
C
AURKB
CNOT3
ERK
WT
CNOT3-DM

**Supplementary Figure 7: Phosphorylation of CNOT3-T292/S294 alters interaction of CNOT3 with ERK in mesendodermal cells**.

**A**.Immunoblotting analysis of the indicated proteins from cell extracts prepared from undifferentiated wild-type (WT) and *Cnot3*-DM ESCs. $\alpha$-Tubulin was used as a loading control.

**B.** PLA detection of endogenous interaction between CNOT3 and ERK using a second pair of antibodies (results from the first antibody pair are shown in Figure 7b). WT and *Cnot3*-DM ESCs were induced to differentiate into mesendodermal cells by incubation for 4 days with BMP4 + FGF2. Positive PLA signals are visible as red dots. Nuclei were counterstained with DAPI and staining for Tubulin was used to mark the cell cytoplasm. Boxed areas represent an enlarged cell indicated by the arrows; scale bar: 10 $\mu$m. PLA dots were quantified from randomly chosen fields from at least 50 cells for each biological replicate. Histogram represents average interactions per cell (dots per cell) as well as the nuclear and cytoplasmic distribution. Error bars represent the Mean ± SEM; $n$=2. (P values calculated by unpaired $t$-test, *P<0.05).

**C.** Negative control single PLAs for analysis shown in Figure 6. The control assays were conducted with conducted with anti-Aurora B only, anti-CNOT3 only or anti-ERK only.

#### Supplementary Table 1

List of primers used for qPCR

| | |
|---|---|
| *Oct-4* | F-5′-CCAATCAGCTTGGGCTAGAG-3′<br>R-5′-CTGGGAAAGGTGTCCCTGTA-3′ |
| *Nanog* | F-5′-TACCTCAGCCTCCAGCAGAT-3′<br>R-5′-GCAATGGATGCTGGGATACT-3′ |
| *Sox2* | F-5'-CACAACTCGGAGATCAGCAA-3'<br>R-5'-CTCCGGGAAGCGTGTACTTA-3' |
| *Sox1* | F-5'-CCTCGGATCTCTGGTCAAGT-3'<br>R-5'-TACAGAGCCGGCAGTCATAC-3' |
| *Fgf5* | F-5′-TCTGGATCTCCTTTGCGTTT-3′<br>R-5′-GGGCTTCGAAAGCACATTTA-3′ |
| *Cdx2* | F-5′-TGGTGTACACAGACCATCAGC-3′<br>R-5′-CCTTGGCTCTGCGGTTCT-3′ |
| *Tead4* | F-5’-ATCCTGACGGAGGAAGGCA-3’<br>R-5'-GCTTGATATGGCGTGCGAT-3' |
| *Brachury* | F-5′-CAGCTGTCGGGGAGCCTGG-3′<br>R-5′-TGCTGCCTGTGAGTCATAAC-3′ |
| *Mixl1* | F-5′-AGTTGCTGGAGCTCGTCTTC-3′<br>R-5′-TTCTGGAACCACACCTGGAT-3′ |
| *Gata4* | F-5′-TTCCTCTCCCAGGAACATCAA-3′<br>R-5′-GCTGCACAACTGGGCTCTACTT-3′ |
| *Gata6* | F-5’-ACAGCCCACTTCTGTGTTCCC-3’<br>R-5’-GTGGGTTGGTCACGTGGTACAG-3’ |
| *Sox17* | F-5’- TTCTGTACACTTTAATGAGGCTGTTC-3’<br>R-5’-TTGTGGGAAGTGGGATCAAG-3’ |
| *Pdgfr* | F-5’-CTGGTGCCTGCCTCCTATGAC-3’<br>R-5’-CACGATCGTTTCTCCTGCCTTAT-3’ |
| *Cxcr4* | F-5’-GAGGCCAAGGAAACTGCTG-3’<br>R-5’-GCGGTCACAGATGTACCTGTC-3’ |
| *Mln51 b* | F-5’- ATGACGATGAGGATCGGAAAAAC-3’<br>R-5’-GTCCCCTTGGGTCGGACTTC-3’ |

#### Supplementary Table 2

List of antibodies

| **Antibody** | **Source** | **Catalouge number** |
|---|---|---|
| Rabbit polyclonal anti-H3 | ABCAM | Cat# ab1791 |
| Rabbit polyclonal anti-CNOT3 | Bethyl Laboratories,Inc | Cat# A302-156A |
| Mouse monoclonal anti-CNOT3 | Abnova | Cat# H00004849-M01 |
| Mouse monoclonal anti-ERK1/2 | Santa Cruz Biotechnology | Cat# sc-514302 |
| Rabbit polyclonal anti-ERK1/2 | Cell Signaling Technology | Cat# 9102 |
| Rabbit polyclonal anti-phospho-ERK1/2(T202/Y204) | Cell Signaling Technology | Cat# 9101 |
| Mouse monoclonal anti-Aurora B | Santa Cruz Biotechnology | Cat# sc-393357 |
| Mouse monoclonal anti-Lamin B1 | Santa Cruz Biotechnology | Cat# sc-374015 |
| Mouse monoclonal anti-$\alpha$ Tubulin | Santa Cruz Biotechnology | Cat# sc-5286 |
| Mouse monoclonal anti-OTX2 | Santa Cruz Biotechnology | Cat# sc-514195 |
| Rabbit polyclonal anti-GATA4 | ABCAM | Cat# ab84593 |
| Rabbit polyclonal anti-$\alpha$ SMA | ABCAM | Cat# ab5694 |
| Mouse monoclonal anti-V5 | ABCAM | Cat# ab27671 |
| Goat polyclonal anti-Brachyury | Santa Cruz Biotechnology | Cat# sc-17743 |
| Rabbit monoclonal anti-FoxA2 | ABCAM | Cat# ab108422 |
| Mouse monoclonal anti-Nestin | Santa Cruz Biotechnology | Cat# sc-23927 |
| Mouse monoclonal anti-$\alpha$ Tubulin-FITC | Merck | Cat# F2168 |
| Aanti-rabbit IgG-HRP | Santa Cruz Biotechnology | Cat# sc-2357 |
| Anti-mouse IgG-HRP | Cell Signaling Technology | Cat# 7076 |
| Donkey anti-goat IgG-Alexa fluor 555 | Thermo Fisher Scientific | Cat# A-21432 |
| Donkey anti-rabbit IgG-Alexa fluor 488 | Thermo Fisher Scientific | Cat# A-21206 |
| Goat anti-mouse IgG-Alexa fluor 488 | Thermo Fisher Scientific | Cat# A-11029 |

### Supplementary Table 3A

RNA sequencing: List of genes downregulated in 4-day differentiated mesendoderm cells from *Cnot3*-DM ESCs compared with differentiated wild-type ESCs

| EnsemblID | baseMean | log2FoldChang | lfcSE | stat | pvalue | padj | mgi_symbol | entrezgene | chromosome_name |
|---|---|---|---|---|---|---|---|---|---|
| ENSMUSG00000053838 | 2482.309155 | -1.157018261 | 0.103489383 | -11.1800672 | 5.11E-29 | 6.73E-25 | Nudcd3 | 209586 | 11 |
| ENSMUSG00000026131 | 11067.97424 | -0.717944873 | 0.087807241 | -8.176374302 | 2.93E-16 | 1.93E-12 | Dst | 13518 | 1 |
| ENSMUSG00000051397 | 1268.069576 | -1.345991691 | 0.168254931 | -7.999716172 | 1.25E-15 | 5.48E-12 | Tacstd2 | 56753 | 6 |
| ENSMUSG00000035632 | 1369.952484 | -0.754342986 | 0.112259751 | -6.71962106 | 1.82E-11 | 4.80E-08 | Cnot3 | 232791 | 7 |
| ENSMUSG00000027070 | 15994.48014 | -1.023105892 | 0.156506894 | -6.537129877 | 6.27E-11 | 1.38E-07 | Lrp2 | 14725 | 2 |
| ENSMUSG00000027820 | 1497.443593 | -0.672144989 | 0.108720951 | -6.18229498 | 6.32E-10 | 1.04E-06 | Mme | 17380 | 3 |
| ENSMUSG00000020758 | 551.3270647 | -1.428458406 | 0.239727538 | -5.958674649 | 2.54E-09 | 3.72E-06 | Itgb4 | 192897 | 11 |
| ENSMUSG00000021061 | 2236.735648 | -0.987576917 | 0.167470576 | -5.897017489 | 3.70E-09 | 4.43E-06 | Spnb1 | 20741 | 12 |
| ENSMUSG00000021182 | 1135.227706 | -0.889362926 | 0.151228429 | -5.880924183 | 4.08E-09 | 4.48E-06 | Ccdc88c | 68339 | 12 |
| ENSMUSG00000033227 | 256.1050069 | -1.666444604 | 0.288426601 | -5.777707739 | 7.57E-09 | 7.13E-06 | Wnt6 | 22420 | 1 |
| ENSMUSG00000058550 | 274.3457105 | -1.267724031 | 0.220558365 | -5.747793938 | 9.04E-09 | 7.94E-06 | Dppa4 | 73693 | 16 |
| ENSMUSG00000068876 | 3552.371463 | -0.646409155 | 0.114422211 | -5.649332837 | 1.61E-08 | 1.33E-05 | Cgn | 70737 | 3 |
| ENSMUSG00000032890 | 417.2103962 | -1.026493014 | 0.189289181 | -5.422882633 | 5.86E-08 | 3.86E-05 | Rims3 | 242662 | 4 |
| ENSMUSG00000055737 | 1081.672873 | -0.86646144 | 0.160855145 | -5.386594496 | 7.18E-08 | 4.51E-05 | Ghr | 14600 | 15 |
| ENSMUSG00000024558 | 729.6436579 | -1.062973842 | 0.198571592 | -5.353101275 | 8.65E-08 | 4.95E-05 | Mapk4 | 225724 | 18 |
| ENSMUSG00000030849 | 1051.836366 | -0.702726847 | 0.131273988 | -5.353130918 | 8.64E-08 | 4.95E-05 | Fgfr2 | 14183 | 7 |
| ENSMUSG00000028444 | 385.1652111 | -1.268432175 | 0.238459807 | -5.31927033 | 1.04E-07 | 5.72E-05 | Cntfr | 12804 | 4 |
| ENSMUSG00000005533 | 1909.520508 | -0.669193516 | 0.126381111 | -5.295043783 | 1.19E-07 | 6.03E-05 | Igf1r | 16001 | 7 |
| ENSMUSG00000038677 | 1295.940121 | -0.615258047 | 0.117193208 | -5.249946303 | 1.52E-07 | 7.16E-05 | Scube3 | 268935 | 17 |
| ENSMUSG00000020023 | 934.8247378 | -0.858819039 | 0.16566568 | -5.184049203 | 2.17E-07 | 9.54E-05 | Tmcc3 | 319880 | 10 |
| ENSMUSG00000020340 | 3443.521921 | -0.708241996 | 0.13697307 | -5.170666006 | 2.33E-07 | 9.92E-05 | Cyfip2 | 76884 | 11 |
| ENSMUSG00000031351 | 853.1260034 | -0.633014664 | 0.125310002 | -5.051589292 | 4.38E-07 | 0.000164957 | Zfp185 | 22673 | X |
| ENSMUSG00000037995 | 2416.855513 | -0.81685748 | 0.161987049 | -5.042733249 | 4.59E-07 | 0.000167981 | Igsf9 | 93842 | 1 |
| ENSMUSG00000026994 | 880.0719507 | -0.688305772 | 0.138261049 | -4.978305742 | 6.41E-07 | 0.000216722 | Galnt3 | 14425 | 2 |
| ENSMUSG00000032087 | 171.7952503 | -1.303279121 | 0.26299458 | -4.955536049 | 7.21E-07 | 0.000237576 | Dscaml1 | 114873 | 9 |
| ENSMUSG00000040728 | 2156.225132 | -0.686425559 | 0.13865033 | -4.950767593 | 7.39E-07 | 0.000237576 | Esrp1 | 207920 | 4 |
| ENSMUSG00000033060 | 1011.747453 | -0.823982468 | 0.169364573 | -4.865140647 | 1.14E-06 | 0.000331632 | Lmo7 | 380928 | 14 |
| ENSMUSG00000025278 | 28303.41362 | -0.638489462 | 0.133540385 | -4.781246229 | 1.74E-06 | 0.00043043 | Flnb | 286940 | 14 |
| ENSMUSG00000060012 | 870.7055147 | -0.769856835 | 0.161255018 | -4.774157383 | 1.80E-06 | 0.000432352 | Kif13b | 16554 | 14 |
| ENSMUSG00000019124 | 167.3783846 | -1.12622219 | 0.236238816 | -4.767303728 | 1.87E-06 | 0.000433672 | Scrn1 | 69938 | 6 |
| ENSMUSG00000031555 | 2010.162034 | -0.606640014 | 0.127915057 | -4.742522346 | 2.11E-06 | 0.000455953 | Adam9 | 11502 | 8 |
| ENSMUSG00000052105 | 2114.618465 | -0.719708416 | 0.151746754 | -4.742825767 | 2.11E-06 | 0.000455953 | 1110012J17Rik | 68617 | 17 |
| ENSMUSG00000063531 | 1705.081337 | -0.773504067 | 0.16296105 | -4.746557937 | 2.07E-06 | 0.000455953 | Sema3e | 20349 | 5 |
| ENSMUSG00000069806 | 565.4159924 | -1.23287421 | 0.259591515 | -4.749285471 | 2.04E-06 | 0.000455953 | Cacng7 | 81904 | 7 |
| ENSMUSG00000027356 | 327.6499736 | -0.936006175 | 0.199234238 | -4.698018697 | 2.63E-06 | 0.000549455 | Fermt1 | 241639 | 2 |
| ENSMUSG00000069045 | 965.6469682 | -0.666732297 | 0.141907633 | -4.698354022 | 2.62E-06 | 0.000549455 | Ddx3y | 26900 | Y |
| ENSMUSG00000034640 | 1466.172569 | -0.771791175 | 0.16472797 | -4.685246667 | 2.80E-06 | 0.000575717 | Tiparp | 99929 | 3 |
| ENSMUSG00000020689 | 438.1493335 | -1.047575528 | 0.228873833 | -4.577087352 | 4.71E-06 | 0.000927296 | Itgb3 | 16416 | 11 |
| ENSMUSG00000031714 | 2662.888399 | -0.620949226 | 0.13615889 | -4.560475093 | 5.10E-06 | 0.000989012 | Gab1 | 14388 | 8 |
| ENSMUSG00000038576 | 971.8499366 | -0.734149212 | 0.161145912 | -4.55580414 | 5.22E-06 | 0.000996593 | Susd4 | 96935 | 1 |
| ENSMUSG00000027500 | 2474.568492 | -0.760402758 | 0.167625832 | -4.536310117 | 5.72E-06 | 0.001047699 | Stmn2 | 20257 | 3 |
| ENSMUSG00000014602 | 6347.665822 | -0.672046889 | 0.14830823 | -4.531420058 | 5.86E-06 | 0.001050004 | Kif1a | 16560 | 1 |
| ENSMUSG00000022197 | 536.7161592 | -1.135428358 | 0.250790471 | -4.527398318 | 5.97E-06 | 0.001050004 | Pdzd2 | 68070 | 15 |
| ENSMUSG00000021838 | 950.7388304 | -0.72783968 | 0.163150473 | -4.461155797 | 8.15E-06 | 0.001249039 | Samd4 | 74480 | 14 |
| ENSMUSG00000038894 | 2713.782975 | -0.593575131 | 0.133334927 | -4.451760275 | 8.52E-06 | 0.001275312 | Irs2 | 384783 | 8 |
| ENSMUSG00000026475 | 506.5810476 | -0.857809037 | 0.194251907 | -4.415961979 | 1.01E-05 | 0.001440329 | Rgs16 | 19734 | 1 |
| ENSMUSG00000024087 | 196.6400177 | -1.057919511 | 0.240086239 | -4.406414617 | 1.05E-05 | 0.001473241 | Cyp1b1 | 13078 | 17 |
| ENSMUSG00000063450 | 780.7824145 | -0.80544092 | 0.183333086 | -4.393320029 | 1.12E-05 | 0.001516477 | Syne2 | 319565 | 12 |
| ENSMUSG00000018849 | 2406.894801 | -0.618289545 | 0.140949816 | -4.386593493 | 1.15E-05 | 0.001548159 | Wwc1 | 211652 | 11 |
| ENSMUSG00000003134 | 545.792238 | -0.806564009 | 0.184049905 | -4.382311474 | 1.17E-05 | 0.001562962 | Tbc1d8 | 54610 | 1 |
| ENSMUSG00000030987 | 1489.503023 | -0.617613927 | 0.141300326 | -4.37093066 | 1.24E-05 | 0.001614092 | Stim1 | 20866 | 7 |
| ENSMUSG00000002799 | 1009.798485 | -0.902324792 | 0.206897161 | -4.361223659 | 1.29E-05 | 0.001631089 | Jag2 | 16450 | 12 |
| ENSMUSG00000019894 | 563.8510355 | -0.996412523 | 0.228527023 | -4.360151859 | 1.30E-05 | 0.001631089 | Slc6a15 | 103098 | 10 |
| ENSMUSG00000025964 | 615.9340152 | -1.14029745 | 0.261821774 | -4.355243006 | 1.33E-05 | 0.001652338 | Adam23 | 23792 | 1 |
| ENSMUSG00000035969 | 1103.178194 | -0.840883566 | 0.195231645 | -4.307106897 | 1.65E-05 | 0.001999564 | Rusc2 | 100213 | 4 |
| ENSMUSG00000043388 | 249.7067451 | -0.859913321 | 0.19982516 | -4.303328567 | 1.68E-05 | 0.002015495 | Tmem130 | 243339 | 5 |
| ENSMUSG00000054452 | 1401.893304 | -0.66899182 | 0.156862142 | -4.264839248 | 2.00E-05 | 0.002353579 | Aes | 14797 | 10 |
| ENSMUSG00000047986 | 679.8603921 | -0.723217059 | 0.169864064 | -4.257622479 | 2.07E-05 | 0.002409321 | Palm3 | 74337 | 8 |
| ENSMUSG00000051554 | 695.1761078 | -0.607316475 | 0.143730516 | -4.225382958 | 2.39E-05 | 0.002686472 | Gm9853 | NA | 8 |
| ENSMUSG00000028337 | 359.330846 | -1.021700227 | 0.242006778 | -4.221783523 | 2.42E-05 | 0.00270661 | Coro2a | 107684 | 4 |
| ENSMUSG00000051375 | 967.1463979 | -0.712646204 | 0.1707267 | -4.174193053 | 2.99E-05 | 0.003244822 | Pcdh1 | 75599 | 18 |
| ENSMUSG00000024462 | 674.5916187 | -0.645406109 | 0.155782016 | -4.143007814 | 3.43E-05 | 0.003556548 | Gabbr1 | 54393 | 17 |
| ENSMUSG00000024998 | 175.4173074 | -1.039560284 | 0.252840762 | -4.111521713 | 3.93E-05 | 0.003863474 | Plce1 | 74055 | 19 |
| ENSMUSG00000043857 | 226.4472698 | -0.965228876 | 0.234854161 | -4.109907492 | 3.96E-05 | 0.003863474 | Mgat5b | 268510 | 11 |
| ENSMUSG00000037860 | 103.9223025 | -1.767514188 | 0.430391727 | -4.106756883 | 4.01E-05 | 0.00388773 | Aim2 | 383619 | 1 |
| ENSMUSG00000063972 | 2760.551781 | -0.669917041 | 0.163954442 | -4.08599506 | 4.39E-05 | 0.004160549 | Nr6a1 | 14536 | 2 |
| ENSMUSG00000019851 | 2176.298047 | -0.58876614 | 0.144669479 | -4.069732905 | 4.71E-05 | 0.004398601 | Perp | 64058 | 10 |
| ENSMUSG00000020467 | 533.7515637 | -0.642716971 | 0.159622143 | -4.026490065 | 5.66E-05 | 0.005040704 | Efemp1 | 216616 | 11 |
| ENSMUSG00000027171 | 736.1395379 | -0.878354678 | 0.218510077 | -4.019744497 | 5.83E-05 | 0.005099059 | Prrg4 | 228413 | 2 |
| ENSMUSG00000028412 | 2407.111743 | -0.666148379 | 0.165826744 | -4.01713476 | 5.89E-05 | 0.005099059 | Slc44a1 | 100434 | 4 |
| ENSMUSG00000018166 | 2057.274688 | -0.775810034 | 0.19395593 | -3.999929433 | 6.34E-05 | 0.005285231 | Erbb3 | 13867 | 10 |
| ENSMUSG00000034275 | 259.8939048 | -0.829067627 | 0.208882964 | -3.969053348 | 7.22E-05 | 0.005869355 | Igsf9b | 235086 | 9 |
| ENSMUSG00000032735 | 60.9293543 | -1.713530065 | 0.433696632 | -3.950987713 | 7.78E-05 | 0.006154372 | Ablim3 | 319713 | 18 |
| ENSMUSG00000059713 | 536.6261686 | -0.779744854 | 0.197399062 | -3.950094027 | 7.81E-05 | 0.006154372 | Rcan3 | 53902 | 4 |
| ENSMUSG00000056608 | 853.0200053 | -0.687715046 | 0.174606548 | -3.938655539 | 8.19E-05 | 0.006314131 | Chd9 | 109151 | 8 |

| | | | | | | | | | |
|---|---|---|---|---|---|---|---|---|---|
| ENSMUSG00000032363 | 306.8802662 | -0.755066597 | 0.192708221 | -3.91818571 | 8.92E-05 | 0.006774951 | Adamts7 | 108153 | 9 |
| ENSMUSG00000044279 | 992.0536055 | -0.643750759 | 0.164325804 | -3.917526916 | 8.95E-05 | 0.006774951 | Crb3 | 224912 | 17 |
| ENSMUSG00000024544 | 132.3469829 | -1.071625592 | 0.273919661 | -3.9121894 | 9.15E-05 | 0.006847782 | D18Ertd653e | 52662 | 18 |
| ENSMUSG00000064043 | 561.5700864 | -0.700442376 | 0.181080778 | -3.868121086 | 0.000109677 | 0.007854437 | Trerf1 | 224829 | 17 |
| ENSMUSG00000046329 | 665.4264395 | -0.749739986 | 0.194180655 | -3.86104366 | 0.000112904 | 0.007916315 | Slc25a23 | 66972 | 17 |
| ENSMUSG00000033826 | 496.3531197 | -0.812364539 | 0.213298814 | -3.808575045 | 0.00013977 | 0.009294436 | Dnahc8 | 13417 | 17 |
| ENSMUSG00000042078 | 365.0013455 | -0.829522031 | 0.21870893 | -3.792812811 | 0.00014895 | 0.009621176 | Svop | 68666 | 5 |
| ENSMUSG00000030790 | 158.2585987 | -1.098491613 | 0.290360693 | -3.783196696 | 0.000154827 | 0.009855821 | Adm | 11535 | 7 |
| ENSMUSG00000017009 | 1374.71206 | -0.757035729 | 0.200182926 | -3.781719761 | 0.000155749 | 0.009866826 | Sdc4 | 20971 | 2 |
| ENSMUSG00000046447 | 233.812982 | -0.817692999 | 0.216319928 | -3.780016972 | 0.000156818 | 0.009887016 | Camk2n1 | 66259 | 4 |
| ENSMUSG00000001870 | 437.9258165 | -0.708633541 | 0.187694492 | -3.775462632 | 0.000159711 | 0.010021483 | Ltbp1 | 268977 | 17 |
| ENSMUSG00000027931 | 249.2730371 | -0.872746922 | 0.231814228 | -3.764854863 | 0.000166646 | 0.010407073 | Npr1 | 18160 | 3 |
| ENSMUSG00000049281 | 180.3654356 | -1.239350705 | 0.330300893 | -3.752186962 | 0.000175299 | 0.010766033 | Scn3b | 235281 | 9 |
| ENSMUSG00000022206 | 87.22724761 | -1.829579116 | 0.488230497 | -3.747367536 | 0.0001787 | 0.010801522 | Npr3 | 18162 | 15 |
| ENSMUSG00000038453 | 147.8639665 | -1.148541121 | 0.306461568 | -3.74774928 | 0.000178428 | 0.010801522 | Srcin1 | 56013 | 11 |
| ENSMUSG00000027330 | 569.172035 | -0.792611094 | 0.213164755 | -3.718302753 | 0.000200566 | 0.011847737 | Cdc25b | 12531 | 2 |
| ENSMUSG00000002228 | 227.0677094 | -0.961737216 | 0.26041732 | -3.693061638 | 0.00022157 | 0.012494774 | Ppm1j | 71887 | 3 |
| ENSMUSG00000017737 | 547.2662676 | -0.916496152 | 0.248191273 | -3.692700964 | 0.000221885 | 0.012494774 | Mmp9 | 17395 | 2 |
| ENSMUSG00000027536 | 485.6112117 | -0.631446152 | 0.171056774 | -3.691441958 | 0.000222986 | 0.012503363 | Chmp4c | 66371 | 3 |
| ENSMUSG00000040274 | 390.2504138 | -0.657073286 | 0.178068502 | -3.690002891 | 0.000224252 | 0.012521026 | Cdk6 | 12571 | 5 |
| ENSMUSG00000051455 | 134.9582589 | -0.940910356 | 0.255694969 | -3.679815676 | 0.000233403 | 0.012922462 | Gm1564 | 268491 | 11 |
| ENSMUSG00000056602 | 274.0242929 | -0.706148426 | 0.193266208 | -3.653760442 | 0.000258427 | 0.013956139 | Fry | 320365 | 5 |
| ENSMUSG00000034593 | 1834.62353 | -0.599763067 | 0.164417084 | -3.647814761 | 0.00026448 | 0.014052644 | Myo5a | 17918 | 9 |
| ENSMUSG00000053559 | 264.8760246 | -0.87769918 | 0.240919945 | -3.643115479 | 0.000269358 | 0.014254335 | Smagp | 207818 | 15 |
| ENSMUSG00000000308 | 435.7158043 | -0.883801225 | 0.243203177 | -3.634003616 | 0.000279057 | 0.014649934 | Ckmt1 | 12716 | 2 |
| ENSMUSG00000039137 | 591.311762 | -0.657287931 | 0.181182373 | -3.62776974 | 0.00028588 | 0.014889491 | Whrn | 73750 | 4 |
| ENSMUSG00000032878 | 148.4474587 | -0.935040212 | 0.259084382 | -3.609018055 | 0.000307358 | 0.015820544 | Ccdc85a | 216613 | 11 |
| ENSMUSG00000019796 | 766.1803887 | -0.632285722 | 0.175370027 | -3.605437782 | 0.000311627 | 0.015915934 | Lrp11 | 237253 | 10 |
| ENSMUSG00000021457 | 209.2378261 | -1.05299028 | 0.291994231 | -3.606202339 | 0.000310711 | 0.015915934 | Syk | 20963 | 13 |
| ENSMUSG00000044976 | 247.7723701 | -0.948932459 | 0.263811167 | -3.597013992 | 0.000321891 | 0.016313692 | Wdr72 | 546144 | 9 |
| ENSMUSG00000039601 | 211.7875114 | -1.092139341 | 0.304249505 | -3.589617479 | 0.000331164 | 0.016592176 | Rcan2 | 53901 | 17 |
| ENSMUSG00000026880 | 461.2632667 | -0.82873851 | 0.232524446 | -3.564091966 | 0.000365118 | 0.017368805 | Stom | 13830 | 2 |
| ENSMUSG00000040488 | 837.778164 | -0.809664275 | 0.227465039 | -3.559510859 | 0.000371546 | 0.017592759 | Ltbp4 | 108075 | 7 |
| ENSMUSG00000020020 | 143.9897749 | -1.125683025 | 0.316725345 | -3.554129925 | 0.000379232 | 0.017669951 | Usp44 | 327799 | 10 |
| ENSMUSG00000042195 | 1106.085107 | -0.682537728 | 0.192144199 | -3.552216165 | 0.000382001 | 0.017724035 | Slc35f2 | 72022 | 9 |
| ENSMUSG00000029426 | 805.7365456 | -0.599074078 | 0.168825938 | -3.548471796 | 0.000387473 | 0.017914871 | Scarb2 | 12492 | 5 |
| ENSMUSG00000027381 | 1222.559113 | -0.642446121 | 0.182407785 | -3.522032357 | 0.000428252 | 0.019325599 | Bcl2l11 | 12125 | 2 |
| ENSMUSG00000034855 | 346.3556108 | -0.747186606 | 0.212318964 | -3.519170368 | 0.000432899 | 0.019374865 | Cxcl10 | 15945 | 5 |
| ENSMUSG00000023232 | 140.1410526 | -1.017096121 | 0.29118083 | -3.493005091 | 0.000477617 | 0.02077084 | Serinc2 | 230779 | 4 |
| ENSMUSG00000021822 | 460.072347 | -0.682484879 | 0.196795947 | -3.467982393 | 0.000524382 | 0.022434341 | Plau | 18792 | 14 |
| ENSMUSG00000040669 | 3905.628905 | -0.596426091 | 0.173603367 | -3.435567536 | 0.000591314 | 0.02427335 | Phc1 | 13619 | 6 |
| ENSMUSG00000029151 | 136.2532322 | -1.286979216 | 0.37696161 | -3.414085633 | 0.000639965 | 0.02540004 | Slc30a3 | 22784 | 5 |
| ENSMUSG00000046314 | 323.5852831 | -0.597445954 | 0.175156452 | -3.410927472 | 0.000647423 | 0.025542198 | Stxbp6 | 217517 | 12 |
| ENSMUSG00000028583 | 1106.222368 | -0.618995315 | 0.181737601 | -3.40598374 | 0.000659261 | 0.025701429 | Pdpn | 14726 | 4 |
| ENSMUSG00000015094 | 435.8115648 | -0.601991886 | 0.177201287 | -3.397220735 | 0.00068074 | 0.026382691 | Npdc1 | 18146 | 2 |
| ENSMUSG00000074796 | 515.7895946 | -0.818326069 | 0.241660001 | -3.386270239 | 0.000708496 | 0.026904463 | Slc4a11 | 269356 | 2 |
| ENSMUSG00000019864 | 523.1608831 | -0.617201861 | 0.183421566 | -3.36493616 | 0.000765614 | 0.027868762 | Rtn4ip1 | 170728 | 10 |
| ENSMUSG00000003282 | 371.9772629 | -0.602278117 | 0.180142276 | -3.343346889 | 0.000827744 | 0.029399399 | Plag1 | 56711 | 4 |
| ENSMUSG00000042659 | 910.8963245 | -0.779017285 | 0.233357962 | -3.338293149 | 0.000842948 | 0.029623913 | Arrdc4 | 66412 | 7 |
| ENSMUSG00000074923 | 140.1519969 | -1.109560024 | 0.335239856 | -3.309749738 | 0.000933794 | 0.031550273 | Pak6 | 214230 | 2 |
| ENSMUSG00000042978 | 766.2424091 | -0.641193459 | 0.193772785 | -3.308996454 | 0.00093631 | 0.031554373 | Sbk1 | 104175 | 7 |
| ENSMUSG00000020583 | 234.7205681 | -1.025261674 | 0.312091576 | -3.285130882 | 0.00101935 | 0.033247466 | Matn3 | 17182 | 12 |
| ENSMUSG00000048915 | 269.4119682 | -0.606089225 | 0.184902421 | -3.277886908 | 0.001045873 | 0.033861098 | Efna5 | 13640 | 17 |
| ENSMUSG00000057329 | 161.7421936 | -0.92519922 | 0.282969476 | -3.269607855 | 0.001076967 | 0.034528444 | Bcl2 | 12043 | 1 |
| ENSMUSG00000027238 | 219.8192346 | -1.035001357 | 0.317780842 | -3.256965867 | 0.0011261 | 0.03566975 | Frmd5 | 228564 | 2 |
| ENSMUSG00000040152 | 12933.2347 | -0.914793522 | 0.281635764 | -3.248144019 | 0.001161605 | 0.036206822 | Thbs1 | 21825 | 2 |
| ENSMUSG00000047793 | 181.0622527 | -0.910896221 | 0.281189357 | -3.239440609 | 0.001197644 | 0.036855036 | Sned1 | 208777 | 1 |
| ENSMUSG00000056476 | 523.7232778 | -0.589617918 | 0.182059933 | -3.238592408 | 0.001201211 | 0.036855036 | Med12l | 329650 | 3 |
| ENSMUSG00000018554 | 312.8843707 | -0.850302513 | 0.262692272 | -3.236876771 | 0.001208456 | 0.03687614 | Ybx2 | 53422 | 11 |
| ENSMUSG00000024883 | 352.7993191 | -0.708711081 | 0.219011592 | -3.235952374 | 0.001212376 | 0.036894873 | Rin1 | 225870 | 19 |
| ENSMUSG00000039238 | 143.5878287 | -0.98972657 | 0.307157331 | -3.22221373 | 0.001272042 | 0.037836791 | Zfp750 | 319530 | 11 |
| ENSMUSG00000036225 | 257.6241029 | -0.711846438 | 0.221128006 | -3.219160021 | 0.001285667 | 0.038155938 | Kctd1 | 106931 | 18 |
| ENSMUSG00000031284 | 160.7919775 | -0.889432342 | 0.277167235 | -3.209009692 | 0.00133193 | 0.039175986 | Pak3 | 18481 | X |
| ENSMUSG00000032251 | 503.1594631 | -0.668634313 | 0.208496249 | -3.206936884 | 0.001341564 | 0.039238495 | Irak1bp1 | 65099 | 9 |
| ENSMUSG00000028789 | 608.8495129 | -0.678086113 | 0.211735774 | -3.202510843 | 0.001362352 | 0.039256702 | Adc | 242669 | 4 |
| ENSMUSG00000015944 | 525.6287015 | -0.751380015 | 0.234709847 | -3.201314416 | 0.001368022 | 0.03927325 | Gatsl2 | 80909 | 5 |
| ENSMUSG00000029314 | 483.1082236 | -0.632702441 | 0.197701589 | -3.200290112 | 0.001372893 | 0.039327421 | Agpat9 | 231510 | 5 |
| ENSMUSG00000043415 | 431.8541099 | -0.694380959 | 0.21709059 | -3.198576961 | 0.001381077 | 0.039395078 | Otud1 | 71198 | 2 |
| ENSMUSG00000024302 | 112.2464352 | -1.375191189 | 0.43247554 | -3.179812639 | 0.001473703 | 0.041141917 | Dtna | 13527 | 18 |
| ENSMUSG00000061451 | 96.30824115 | -1.302076757 | 0.410152798 | -3.174613857 | 0.00150036 | 0.041447056 | Tmem151a | 381199 | 19 |
| ENSMUSG00000019256 | 537.2204281 | -0.641109308 | 0.202945557 | -3.159021159 | 0.001583 | 0.042832014 | Ahr | 11622 | 12 |
| ENSMUSG00000041703 | 786.1848823 | -0.692415194 | 0.219274804 | -3.157750826 | 0.001589914 | 0.042924203 | Zic5 | 65100 | 14 |
| ENSMUSG00000026463 | 370.6140132 | -0.825209385 | 0.261564195 | -3.154901933 | 0.001605521 | 0.043175398 | Atp2b4 | 381290 | 1 |
| ENSMUSG00000037846 | 268.9204955 | -0.85917001 | 0.273521703 | -3.14114017 | 0.001682915 | 0.044529652 | Rtkn2 | 170799 | 10 |
| ENSMUSG00000069170 | 89.53057658 | -1.195016711 | 0.381363337 | -3.133538535 | 0.001727122 | 0.045038854 | Gpr98 | 110789 | 13 |
| ENSMUSG00000015312 | 3254.58288 | -0.601931231 | 0.192217506 | -3.131510981 | 0.001739093 | 0.045110288 | Gadd45b | 17873 | 10 |
| ENSMUSG00000022101 | 95.66484114 | -0.925418559 | 0.296748311 | -3.118530161 | 0.001817555 | 0.046504713 | Fgf17 | 14171 | 14 |
| ENSMUSG00000034656 | 312.0232071 | -0.629091519 | 0.202390855 | -3.108300122 | 0.001881669 | 0.047774084 | Cacna1a | 12286 | 8 |
| ENSMUSG00000032625 | 303.023047 | -0.827390575 | 0.266676388 | -3.102601557 | 0.001918277 | 0.048138771 | Thsd7a | 330267 | 6 |
| ENSMUSG00000085328 | 1123.856096 | -0.627964356 | 0.202432769 | -3.10208845 | 0.001921605 | 0.048138771 | Gm17131 | NA | 5 |

### Supplementary Table 3B

RNA sequencing: List of genes upregulated in 4-day differentiated mesendoderm cells from *Cnot3*-DM ESCs compared with differentiated wild-type ESCs

| EnsemblID | baseMean | log2FoldChange | lfcSE | stat | pvalue | padj | mgi_symbol | entrezgene | chromosome_name |
|---|---|---|---|---|---|---|---|---|---|
| ENSMUSG00000007097 | 283.3887789 | 1.449134136 | 0.200865927 | 7.214434828 | 5.42E-13 | 1.78E-09 | Atp1a2 | 98660 | 1 |
| ENSMUSG00000086266 | 242.2075032 | 1.826247063 | 0.285839193 | 6.389071569 | 1.67E-10 | 3.14E-07 | Igf2as | NA | 7 |
| ENSMUSG00000063229 | 14267.07335 | 0.730361982 | 0.123370722 | 5.920059245 | 3.22E-09 | 4.24E-06 | Ldha | 16828 | 7 |
| ENSMUSG00000052396 | 532.129582 | 0.972038727 | 0.167788707 | 5.793230945 | 6.90E-09 | 7.00E-06 | Mogat2 | 233549 | 7 |
| ENSMUSG00000040856 | 182.8880384 | 1.950298475 | 0.349569919 | 5.579137013 | 2.42E-08 | 1.81E-05 | Dlk1 | 13386 | 12 |
| ENSMUSG00000061723 | 77.28008179 | 2.8924795 | 0.545377262 | 5.303630539 | 1.14E-07 | 5.98E-05 | Tnnt3 | 21957 | 7 |
| ENSMUSG00000021403 | 239.9507425 | 1.621103129 | 0.310118623 | 5.227364661 | 1.72E-07 | 7.81E-05 | Serpinb9b | 20706 | 13 |
| ENSMUSG00000007872 | 1876.535247 | 0.892355993 | 0.173750694 | 5.135841321 | 2.81E-07 | 0.000112158 | Id3 | 15903 | 4 |
| ENSMUSG00000048528 | 47.44919983 | 3.515357332 | 0.683742816 | 5.141344446 | 2.73E-07 | 0.000112158 | Nkx1-2 | 20231 | 7 |
| ENSMUSG00000042745 | 1672.038251 | 0.848185828 | 0.167880816 | 5.05230942 | 4.36E-07 | 0.000164957 | Id1 | 15901 | 2 |
| ENSMUSG00000035960 | 2107.286392 | 0.711616606 | 0.141882113 | 5.015548401 | 5.29E-07 | 0.000188333 | Apex1 | 11792 | 14 |
| ENSMUSG00000024565 | 171.6114202 | 1.974201467 | 0.396227498 | 4.982494844 | 6.28E-07 | 0.000216722 | Sall3 | 20689 | 18 |
| ENSMUSG00000024232 | 752.2929805 | 1.153132173 | 0.235330684 | 4.900050228 | 9.58E-07 | 0.000300599 | Bambi | 68010 | 18 |
| ENSMUSG00000019846 | 341.7072104 | 1.067727329 | 0.218291102 | 4.891300292 | 1.00E-06 | 0.000306969 | Lama4 | 16775 | 10 |
| ENSMUSG00000030041 | 156.1775192 | 1.165738862 | 0.238972297 | 4.878133896 | 1.07E-06 | 0.000320723 | D6Mm5e | 110958 | 6 |
| ENSMUSG00000011256 | 1238.676473 | 0.638469088 | 0.131298154 | 4.862742318 | 1.16E-06 | 0.000331632 | Adam19 | 11492 | 11 |
| ENSMUSG00000021613 | 31.82754114 | 2.978984914 | 0.614289898 | 4.849477297 | 1.24E-06 | 0.00034261 | Hapln1 | 12950 | 13 |
| ENSMUSG00000027009 | 59.42934805 | 2.673006928 | 0.551379214 | 4.847855818 | 1.25E-06 | 0.00034261 | Itga4 | 16401 | 2 |
| ENSMUSG00000039239 | 1388.027084 | 0.778529949 | 0.161102813 | 4.832503745 | 1.35E-06 | 0.000351685 | Tgfb2 | 21808 | 1 |
| ENSMUSG00000043342 | 60.05455736 | 2.405457026 | 0.497961359 | 4.830609811 | 1.36E-06 | 0.000351685 | Hoxd9 | 15438 | 2 |
| ENSMUSG00000042417 | 109.6058058 | 1.979187385 | 0.414164643 | 4.778745404 | 1.76E-06 | 0.00043043 | Ccno | 218630 | 13 |
| ENSMUSG00000015053 | 126.4464755 | 1.563911628 | 0.328115223 | 4.766348894 | 1.88E-06 | 0.000433672 | Gata2 | 14461 | 6 |
| ENSMUSG00000020388 | 202.5452985 | 1.646249561 | 0.353130359 | 4.661874915 | 3.13E-06 | 0.000635216 | Pdlim4 | 30794 | 11 |
| ENSMUSG00000022464 | 7319.351693 | 0.695393563 | 0.153115136 | 4.541638275 | 5.58E-06 | 0.001047699 | Slc38a4 | 69354 | 15 |
| ENSMUSG00000030022 | 653.6365516 | 0.820953259 | 0.181391176 | 4.525872085 | 6.01E-06 | 0.001050004 | Adamts9 | 101401 | 6 |
| ENSMUSG00000032221 | 259.278792 | 1.200599853 | 0.265359727 | 4.524423759 | 6.06E-06 | 0.001050004 | Mns1 | 17427 | 9 |
| ENSMUSG00000051851 | 174.4324088 | 1.693380282 | 0.374948169 | 4.516304978 | 6.29E-06 | 0.001076886 | Cxx1c | 72865 | X |
| ENSMUSG00000006386 | 51.51781704 | 1.952076191 | 0.434110779 | 4.496723617 | 6.90E-06 | 0.001165803 | Tek | 21687 | 4 |
| ENSMUSG00000026728 | 6644.127629 | 0.876829377 | 0.195192869 | 4.492117883 | 7.05E-06 | 0.001176229 | Vim | 22352 | 2 |
| ENSMUSG00000028766 | 230.0251879 | 1.408415984 | 0.313732949 | 4.489219218 | 7.15E-06 | 0.001177442 | Alpl | 11647 | 4 |
| ENSMUSG00000036377 | 893.439453 | 0.98638567 | 0.21989239 | 4.485765384 | 7.27E-06 | 0.001181906 | C530008M17Rik | 320827 | 5 |
| ENSMUSG00000033880 | 130.3966892 | 1.339929887 | 0.299409948 | 4.475235028 | 7.63E-06 | 0.001217489 | Lgals3bp | 19039 | 11 |
| ENSMUSG00000019235 | 107.5336986 | 1.462818247 | 0.327422718 | 4.467674862 | 7.91E-06 | 0.001225839 | Rps6kl1 | 238323 | 12 |
| ENSMUSG00000085795 | 386.3765089 | 0.884719146 | 0.198607832 | 4.454603505 | 8.40E-06 | 0.001272995 | Zfp703 | 353310 | 8 |
| ENSMUSG00000039231 | 1360.553905 | 0.676627841 | 0.152485794 | 4.437317224 | 9.11E-06 | 0.001333615 | Suv39h1 | 20937 | X |
| ENSMUSG00000031073 | 146.312395 | 2.320222907 | 0.524376388 | 4.424728039 | 9.66E-06 | 0.001398266 | Fgf15 | 14170 | 7 |
| ENSMUSG00000061082 | 102.4369856 | 2.178612466 | 0.495416156 | 4.397540211 | 1.09E-05 | 0.00150279 | Plac1 | 56096 | X |
| ENSMUSG00000042436 | 67.17036567 | 1.876224146 | 0.428404791 | 4.379559205 | 1.19E-05 | 0.001567004 | Mfap4 | 76293 | 11 |
| ENSMUSG00000043445 | 518.6094127 | 0.677682134 | 0.155357573 | 4.362079821 | 1.29E-05 | 0.001631089 | Pgp | 67078 | 17 |
| ENSMUSG00000025665 | 3751.887593 | 0.805315731 | 0.185737593 | 4.335771343 | 1.45E-05 | 0.001788745 | Rps6ka6 | 67071 | X |
| ENSMUSG00000070348 | 3039.659541 | 0.700859532 | 0.163380147 | 4.289747231 | 1.79E-05 | 0.002123475 | Ccnd1 | 12443 | 7 |
| ENSMUSG00000037664 | 1709.113048 | 0.736383655 | 0.173329578 | 4.248459278 | 2.15E-05 | 0.002487977 | Cdkn1c | 12577 | 7 |
| ENSMUSG00000025969 | 1081.150516 | 0.921039795 | 0.217007418 | 4.244277936 | 2.19E-05 | 0.002512778 | Nrp2 | 18187 | 1 |
| ENSMUSG00000031891 | 20.72393859 | 3.045214588 | 0.7195039 | 4.232380935 | 2.31E-05 | 0.002626657 | Hsd11b2 | 15484 | 8 |
| ENSMUSG00000001657 | 19.15229534 | 3.781989211 | 0.906268359 | 4.173144936 | 3.00E-05 | 0.003244822 | Hoxc8 | 15426 | 15 |
| ENSMUSG00000027669 | 259.899371 | 1.046158548 | 0.250495145 | 4.176362572 | 2.96E-05 | 0.003244822 | Gnb4 | 14696 | 3 |
| ENSMUSG00000025776 | 117.1059428 | 1.575001062 | 0.377629575 | 4.170756652 | 3.04E-05 | 0.003247022 | Crispld1 | 83691 | 1 |
| ENSMUSG00000031074 | 96.12088287 | 2.14042192 | 0.513378556 | 4.169285789 | 3.06E-05 | 0.003247022 | Fgf3 | 14174 | 7 |
| ENSMUSG00000023484 | 217.609032 | 1.067365499 | 0.256237514 | 4.165531754 | 3.11E-05 | 0.003274513 | Prph | 19132 | 15 |
| ENSMUSG00000029646 | 316.2348609 | 2.510042416 | 0.606294098 | 4.139975015 | 3.47E-05 | 0.00357574 | Cdx2 | 12591 | 5 |
| ENSMUSG00000031963 | 119.5038363 | 1.700607264 | 0.411889804 | 4.128791849 | 3.65E-05 | 0.003696395 | Bmper | 73230 | 9 |
| ENSMUSG00000036523 | 439.6896435 | 0.957871636 | 0.233020854 | 4.110669148 | 3.95E-05 | 0.003863474 | Greb1 | 268527 | 12 |
| ENSMUSG00000028369 | 74.27939726 | 1.9939375 | 0.48661602 | 4.097558278 | 4.18E-05 | 0.003986817 | Svep1 | 64817 | 4 |
| ENSMUSG00000070407 | 298.6815831 | 1.351693942 | 0.333165015 | 4.057130495 | 4.97E-05 | 0.004581984 | Hs3st3b1 | 54710 | 11 |
| ENSMUSG00000005087 | 800.4529198 | 0.606755231 | 0.150575241 | 4.029581665 | 5.59E-05 | 0.005040704 | Cd44 | 12505 | 2 |
| ENSMUSG00000033350 | 96.85296273 | 1.666840896 | 0.414939591 | 4.017068823 | 5.89E-05 | 0.005099059 | Chst2 | 54371 | 9 |
| ENSMUSG00000052776 | 39.68508716 | 2.111008534 | 0.525655601 | 4.015953662 | 5.92E-05 | 0.005099059 | Oas1a | 246730 | 5 |
| ENSMUSG00000031217 | 675.576237 | 0.839185623 | 0.209316302 | 4.009174705 | 6.09E-05 | 0.005213585 | Efnb1 | 13641 | X |
| ENSMUSG00000038507 | 328.3227396 | 1.150761133 | 0.287318155 | 4.005180722 | 6.20E-05 | 0.005252179 | Parp12 | 243771 | 6 |
| ENSMUSG00000031734 | 62.2475798 | 1.549274621 | 0.387329704 | 3.999885898 | 6.34E-05 | 0.005285231 | Irx3 | 16373 | 8 |
| ENSMUSG00000025105 | 43.95377984 | 1.930103703 | 0.485787627 | 3.973142981 | 7.09E-05 | 0.005841565 | Bnc1 | 12173 | 7 |
| ENSMUSG00000040212 | 127.1896627 | 1.203376346 | 0.30317857 | 3.969199895 | 7.21E-05 | 0.005869355 | Emp3 | 13732 | 7 |
| ENSMUSG00000079197 | 2904.588691 | 0.642758069 | 0.162198453 | 3.962787905 | 7.41E-05 | 0.00598863 | Psme2 | 19188 | 14 |
| ENSMUSG00000057615 | 73.76704192 | 1.408899552 | 0.356770098 | 3.949040459 | 7.85E-05 | 0.006154372 | Ldoc1 | 434784 | X |
| ENSMUSG00000031486 | 816.9286127 | 1.566821883 | 0.397424434 | 3.942439745 | 8.07E-05 | 0.00625186 | Gpr124 | 78560 | 8 |
| ENSMUSG00000039316 | 391.4408762 | 1.075879857 | 0.27329916 | 3.936637998 | 8.26E-05 | 0.006330408 | Rftn1 | 76438 | 17 |
| ENSMUSG00000037169 | 1248.113109 | 0.724279625 | 0.185039421 | 3.914190948 | 9.07E-05 | 0.006830041 | Mycn | 18109 | 12 |
| ENSMUSG00000031075 | 47.85274647 | 2.063109925 | 0.532534201 | 3.874136012 | 0.000107004 | 0.007747181 | Ano1 | 101772 | 7 |
| ENSMUSG00000026104 | 300.186188 | 0.986070243 | 0.255395368 | 3.860955862 | 0.000112944 | 0.007916315 | Stat1 | 20846 | 1 |
| ENSMUSG00000062393 | 504.1113151 | 1.008835461 | 0.261997839 | 3.85054879 | 0.000117853 | 0.008216693 | Dgkk | 331374 | X |
| ENSMUSG00000054072 | 56.35084834 | 2.241177558 | 0.583478525 | 3.841062629 | 0.000122503 | 0.008363835 | Iigp1 | 60440 | 18 |
| ENSMUSG00000050335 | 399.0295382 | 0.671277109 | 0.175700922 | 3.82056679 | 0.000133145 | 0.008951307 | Lgals3 | 16854 | 14 |
| ENSMUSG00000025491 | 631.2739523 | 0.854121062 | 0.224268851 | 3.808469427 | 0.00013983 | 0.009294436 | Ifitm1 | 68713 | 7 |
| ENSMUSG00000033585 | 305.3477676 | 0.962534675 | 0.253009146 | 3.804347356 | 0.000142179 | 0.00932083 | Ndn | 17984 | 7 |
| ENSMUSG00000029201 | 1059.029646 | 0.771055882 | 0.202983816 | 3.798607685 | 0.000145511 | 0.009492085 | Ugdh | 22235 | 5 |

| | | | | | | | | | |
|---|---|---|---|---|---|---|---|---|---|
| ENSMUSG00000028005 | 186.1039764 | 1.230694363 | 0.324826933 | 3.788769458 | 0.000151395 | 0.009731396 | Gucy1b3 | 54195 | 3 |
| ENSMUSG00000055653 | 5240.324991 | 0.671074446 | 0.177285211 | 3.785281593 | 0.000153535 | 0.009820999 | Gpc3 | 14734 | X |
| ENSMUSG00000000142 | 609.1554334 | 0.944374794 | 0.251414288 | 3.756249494 | 0.000172479 | 0.010670199 | Axin2 | 12006 | 11 |
| ENSMUSG00000032014 | 74.76513024 | 1.282519972 | 0.343365203 | 3.735148346 | 0.000187605 | 0.011236661 | Oaf | 102644 | 9 |
| ENSMUSG00000003348 | 716.4835036 | 0.616075637 | 0.16649816 | 3.700194855 | 0.000215434 | 0.012306699 | Mob3a | 208228 | 10 |
| ENSMUSG00000024538 | 1170.442418 | 0.71215328 | 0.19248821 | 3.699724155 | 0.000215834 | 0.012306699 | Ppic | 19038 | 18 |
| ENSMUSG00000050105 | 64.76296614 | 1.781500523 | 0.484483418 | 3.677113513 | 0.000235888 | 0.012946424 | Grrp1 | 72690 | 4 |
| ENSMUSG00000074749 | 190.5536996 | 0.78437374 | 0.213368423 | 3.676147238 | 0.000236783 | 0.012946424 | Plk1s1 | 228730 | 2 |
| ENSMUSG00000037347 | 139.9911112 | 1.270076496 | 0.346606631 | 3.66431678 | 0.000248 | 0.013448115 | Chst7 | 60322 | X |
| ENSMUSG00000024659 | 557.8192861 | 0.863236619 | 0.239754998 | 3.600494779 | 0.000317612 | 0.016158981 | Anxa1 | 16952 | 19 |
| ENSMUSG00000020734 | 43.90858602 | 1.771977086 | 0.493508198 | 3.590572745 | 0.000329952 | 0.016592176 | Grin2c | 14813 | 11 |
| ENSMUSG00000026347 | 81.91222271 | 1.2919917 | 0.361841759 | 3.570598665 | 0.000356166 | 0.017254422 | Tmem163 | 72160 | 1 |
| ENSMUSG00000025723 | 84.57747378 | 1.292332265 | 0.362378426 | 3.566250558 | 0.000362125 | 0.01732571 | Nmb | 68039 | 7 |
| ENSMUSG00000000690 | 22.8452417 | 2.976216251 | 0.83650917 | 3.557900329 | 0.000373831 | 0.017592759 | Hoxb6 | 15414 | 11 |
| ENSMUSG00000006200 | 128.0443345 | 1.383350252 | 0.388731125 | 3.558630024 | 0.000372794 | 0.017592759 | Rhox6 | 19202 | X |
| ENSMUSG00000000753 | 88.00235774 | 1.138397562 | 0.320319141 | 3.553947978 | 0.000379494 | 0.017669951 | Serpinf1 | 20317 | 11 |
| ENSMUSG00000048583 | 20924.62298 | 1.763778333 | 0.49791324 | 3.542340695 | 0.000396593 | 0.018217742 | Igf2 | 16002 | 7 |
| ENSMUSG00000019779 | 158.3636336 | 1.204358543 | 0.341072397 | 3.531093558 | 0.000413845 | 0.018804276 | Frk | 14302 | 10 |
| ENSMUSG00000029826 | 572.1774945 | 0.752399816 | 0.213025695 | 3.531967421 | 0.00041248 | 0.018804276 | Zc3hav1 | 78781 | 6 |
| ENSMUSG00000027985 | 410.4341479 | 0.986836582 | 0.280186713 | 3.522067742 | 0.000428195 | 0.019325599 | Lef1 | 16842 | 3 |
| ENSMUSG00000030110 | 48.45422129 | 1.655839402 | 0.470415629 | 3.519949808 | 0.000431628 | 0.019374865 | Ret | 19713 | 6 |
| ENSMUSG00000018819 | 97.32309034 | 1.148317815 | 0.327149214 | 3.51007359 | 0.000447983 | 0.01987565 | Lsp1 | 16985 | 7 |
| ENSMUSG00000027102 | 30.12658109 | 2.033848818 | 0.581120985 | 3.499871574 | 0.000465482 | 0.020445537 | Hoxd8 | 15437 | 2 |
| ENSMUSG00000039081 | 26.98815731 | 2.214462454 | 0.636521309 | 3.47900757 | 0.000503274 | 0.021743103 | Zfp503 | 218820 | 14 |
| ENSMUSG00000052957 | 321.0690726 | 0.867159787 | 0.249229297 | 3.479365378 | 0.000502603 | 0.021743103 | Gas1 | 14451 | 13 |
| ENSMUSG00000026124 | 93.10863363 | 1.807090798 | 0.520759212 | 3.470108176 | 0.000520249 | 0.022361798 | Cfc1 | 12627 | 1 |
| ENSMUSG00000055809 | 212.4537795 | 0.891800405 | 0.257023273 | 3.469726287 | 0.000520989 | 0.022361798 | 6030429G01Rik | 436022 | 7 |
| ENSMUSG00000044408 | 529.9701348 | 0.629857439 | 0.182336132 | 3.454375347 | 0.000551569 | 0.023146581 | 1110002B05Rik | 104725 | 12 |
| ENSMUSG00000042102 | 43.76501694 | 2.103421971 | 0.610299826 | 3.446538705 | 0.000567817 | 0.023528703 | Dmgdh | 74129 | 13 |
| ENSMUSG00000027954 | 234.1608836 | 0.783754634 | 0.228435287 | 3.430970079 | 0.000601427 | 0.024535612 | Efna1 | 13636 | 3 |
| ENSMUSG00000026739 | 249.7036658 | 0.851366901 | 0.248433032 | 3.426947278 | 0.000610408 | 0.02474874 | Bmi1 | 12151 | 2 |
| ENSMUSG00000045333 | 333.6674446 | 0.908403874 | 0.265362393 | 3.423257766 | 0.000618754 | 0.024933694 | Zfp423 | 94187 | 8 |
| ENSMUSG00000028069 | 662.3822588 | 0.651516385 | 0.190449804 | 3.420934915 | 0.000624063 | 0.024994752 | Gpatch4 | 66614 | 3 |
| ENSMUSG00000031661 | 204.2028134 | 1.259915594 | 0.368520495 | 3.418848096 | 0.000628868 | 0.025110899 | Nkd1 | 93960 | 8 |
| ENSMUSG00000072944 | 787.4680846 | 0.635591348 | 0.186127098 | 3.414824355 | 0.000638232 | 0.02540004 | Nup62cl | 279706 | X |
| ENSMUSG00000033774 | 23.57056702 | 2.164246882 | 0.635288071 | 3.406717332 | 0.000657492 | 0.025701429 | Npbwr1 | 226304 | 1 |
| ENSMUSG00000029778 | 77.98337059 | 1.323731271 | 0.39044806 | 3.390287736 | 0.000698193 | 0.026822417 | Adcyap1r1 | 11517 | 6 |
| ENSMUSG00000028212 | 279.8024987 | 0.686629797 | 0.203260622 | 3.378075836 | 0.000729949 | 0.02732541 | Ccne2 | 12448 | 4 |
| ENSMUSG00000042258 | 61.77892304 | 2.059807648 | 0.610147096 | 3.375919775 | 0.000735694 | 0.027357856 | Isl1 | 16392 | 13 |
| ENSMUSG00000033763 | 586.2635763 | 0.591255605 | 0.175488134 | 3.369205613 | 0.000753852 | 0.027714515 | Mtss1l | 244654 | 8 |
| ENSMUSG00000070526 | 50.44535786 | 1.859870292 | 0.552354863 | 3.367165596 | 0.000759451 | 0.027721002 | Peg12 | 27412 | 7 |
| ENSMUSG00000024697 | 27.8346586 | 2.549456864 | 0.758440407 | 3.361446515 | 0.000775354 | 0.028071939 | Gna14 | 14675 | 19 |
| ENSMUSG00000068048 | 64.82582848 | 1.55728285 | 0.463566232 | 3.359353512 | 0.000781251 | 0.028204216 | Rhox9 | 104384 | X |
| ENSMUSG00000023942 | 491.9064894 | 0.588092262 | 0.175418653 | 3.35250699 | 0.000800832 | 0.028753585 | Slc29a1 | 63959 | 17 |
| ENSMUSG00000046352 | 48.39730106 | 1.537451 | 0.461628417 | 3.330494706 | 0.000866918 | 0.030061526 | Gjb2 | 14619 | 14 |
| ENSMUSG00000022346 | 626.0440882 | 0.78145252 | 0.234757534 | 3.32876439 | 0.000872322 | 0.030169508 | Myc | 17869 | 15 |
| ENSMUSG00000040631 | 611.2569082 | 1.563915446 | 0.470624056 | 3.32306737 | 0.000890334 | 0.030551907 | Dok4 | 114255 | 8 |
| ENSMUSG00000029765 | 126.4924966 | 1.448277968 | 0.437119167 | 3.313233729 | 0.000922239 | 0.031401395 | Plxna4 | 243743 | 6 |
| ENSMUSG00000048450 | 523.6718304 | 2.184551489 | 0.659978105 | 3.310036308 | 0.000932839 | 0.031550273 | Msx1 | 17701 | 5 |
| ENSMUSG00000029298 | 92.57453724 | 1.508801165 | 0.457125488 | 3.300627956 | 0.000964687 | 0.032181474 | Gbp9 | 236573 | 5 |
| ENSMUSG00000032897 | 643.2071938 | 0.633624254 | 0.19220442 | 3.296616455 | 0.00097857 | 0.032480155 | Nfyc | 18046 | 4 |
| ENSMUSG00000092035 | 34105.14985 | 0.937321237 | 0.284822182 | 3.29089972 | 0.000998675 | 0.032900921 | Peg10 | 170676 | 6 |
| ENSMUSG00000032291 | 148.6599521 | 0.974592154 | 0.296826366 | 3.283374608 | 0.001025723 | 0.033372709 | Crabp1 | 12903 | 9 |
| ENSMUSG00000000120 | 46.45589628 | 1.920260031 | 0.586139187 | 3.276116105 | 0.001052453 | 0.033990616 | Ngfr | 18053 | 11 |
| ENSMUSG00000019789 | 91.61182862 | 1.103174145 | 0.338106508 | 3.262800681 | 0.001103171 | 0.035112274 | Hey2 | 15214 | 10 |
| ENSMUSG00000030170 | 122.0665686 | 1.417265475 | 0.434509978 | 3.261755882 | 0.001107245 | 0.035157014 | Wnt5b | 22419 | 6 |
| ENSMUSG00000025529 | 132.5336115 | 1.26178139 | 0.388288596 | 3.249596826 | 0.001155687 | 0.036206822 | Zfp711 | 245595 | X |
| ENSMUSG00000036545 | 75.95336569 | 1.247597701 | 0.384035388 | 3.248652962 | 0.001159528 | 0.036206822 | Adamts2 | 216725 | 11 |
| ENSMUSG00000087365 | 159.8753079 | 1.226300099 | 0.377229288 | 3.250808299 | 0.001150774 | 0.036206822 | C430049B03Rik | NA | X |
| ENSMUSG00000079481 | 173.5593355 | 1.203221511 | 0.370674722 | 3.246030657 | 0.001170262 | 0.03628364 | Nhsl2 | 100042480 | X |
| ENSMUSG00000022790 | 116.9654681 | 1.118268383 | 0.346024388 | 3.231761756 | 0.001230296 | 0.037182588 | Igsf11 | 207683 | 16 |
| ENSMUSG00000038244 | 160.2741548 | 0.773104716 | 0.239588898 | 3.226796908 | 0.001251843 | 0.037404836 | Mical2 | 320878 | 7 |
| ENSMUSG00000017493 | 1436.169009 | 1.268390188 | 0.393229761 | 3.225570176 | 0.00125722 | 0.037480519 | Igfbp4 | 16010 | 11 |
| ENSMUSG00000046743 | 40.60609731 | 2.044361415 | 0.635928622 | 3.214765533 | 0.001305511 | 0.038657797 | Fat4 | 329628 | 3 |
| ENSMUSG00000000567 | 124.0173741 | 1.123578986 | 0.351478328 | 3.196723375 | 0.001389982 | 0.039498407 | Sox9 | 20682 | 11 |
| ENSMUSG00000021506 | 37.8116986 | 1.653876652 | 0.517496003 | 3.195921595 | 0.00139385 | 0.039498407 | Pitx1 | 18740 | 13 |
| ENSMUSG00000038193 | 44.17411324 | 2.125041283 | 0.670823095 | 3.167811751 | 0.001535909 | 0.041951108 | Hand2 | 15111 | 8 |
| ENSMUSG00000001864 | 287.8491282 | 0.665475882 | 0.210655465 | 3.159072472 | 0.001582721 | 0.042832014 | Aif1l | 108897 | 2 |
| ENSMUSG00000067786 | 135.1318425 | 1.117083388 | 0.353820947 | 3.157199699 | 0.001592922 | 0.042924203 | Nnat | 18111 | 2 |
| ENSMUSG00000042821 | 371.0679347 | 0.874122045 | 0.277407294 | 3.151042043 | 0.001626891 | 0.043548472 | Snai1 | 20613 | 2 |
| ENSMUSG00000058806 | 27.30566452 | 2.282372194 | 0.724246316 | 3.151375633 | 0.001625034 | 0.043548472 | Col13a1 | 12817 | 10 |
| ENSMUSG00000031871 | 21.8326788 | 2.116838627 | 0.673511093 | 3.142989996 | 0.001672316 | 0.044338233 | Cdh5 | 12562 | 8 |
| ENSMUSG00000022449 | 32.94683304 | 2.094334543 | 0.667836104 | 3.136000781 | 0.001712687 | 0.044894731 | Adamts20 | 223838 | 15 |
| ENSMUSG00000053219 | 68.2590431 | 1.138872646 | 0.363457361 | 3.133442238 | 0.001727689 | 0.045038854 | Raet1e | 379043 | 10 |
| ENSMUSG00000054716 | 246.7132298 | 0.72212711 | 0.231132016 | 3.124305859 | 0.001782251 | 0.045869275 | Zfp771 | 244216 | 7 |
| ENSMUSG00000029544 | 140.2233861 | 1.385115588 | 0.446273032 | 3.10374029 | 0.00191091 | 0.048053549 | Cabp1 | 29867 | 5 |
| ENSMUSG00000049288 | 100.0647123 | 1.174199241 | 0.378287247 | 3.103988438 | 0.001909308 | 0.048053549 | Lix1l | 280411 | 3 |
| ENSMUSG00000021614 | 1019.126802 | 1.2602381 | 0.407274167 | 3.094323678 | 0.001972621 | 0.049043836 | Vcan | 13003 | 13 |
